# SynGAP regulates synaptic plasticity and cognition independent of its catalytic activity

**DOI:** 10.1101/2023.08.06.552111

**Authors:** Yoichi Araki, Kacey E. Rajkovich, Elizabeth E. Gerber, Timothy R. Gamache, Richard C. Johnson, Thanh Hai Tran, Bian Liu, Ingie Hong, Alfredo Kirkwood, Richard Huganir

## Abstract

SynGAP is an abundant synaptic GTPase-activating protein (GAP) critical for synaptic plasticity, learning, memory, and cognition. Mutations in *SYNGAP1* in humans result in intellectual disability, autistic-like behaviors, and epilepsy. Heterozygous *Syngap1* knockout mice display deficits in synaptic plasticity, learning, and memory, and exhibit seizures. It is unclear whether SynGAP imparts structural properties at synapses independent of its GAP activity. Here, we report that inactivating mutations within the SynGAP GAP domain do not inhibit synaptic plasticity or cause behavioral deficits. Instead, SynGAP modulates synaptic strength by physically competing with the AMPA-receptor-TARP complex, the major excitatory receptor complex in the brain, in the formation of molecular condensates with synaptic scaffolding proteins. These results have significant implications for the development of therapeutic treatments for *SYNGAP1*- related neurodevelopmental disorders.

**One-Sentence Summary:** SynGAP regulates synaptic plasticity and cognition due to its phase separation properties instead of its catalytic activity.

## Main Text

Long-term potentiation (LTP) is a major form of synaptic plasticity in the brain, which is thought to underlie learning, memory, and other higher-order brain processes ^1–3^. LTP has been a central focus in neuroscience for decades, and the biochemical signaling cascades underlying LTP have been investigated in great depth. Synaptic potentiation during LTP is mediated by increases in synaptic α-amino-3-hydroxy-5- methyl-4-isoxazolepropionic acid receptors (AMPARs), the major excitatory neurotransmitter receptors in the brain ^1–3^. However, it remains unclear how LTP induction leads to the stable trapping of AMPARs at the synapse to establish and maintain increased synaptic strength. One leading hypothesis involves the diffusional trapping of plasma-membrane-inserted AMPARs by binding to proteinaceous binding “slots” in the postsynaptic density (PSD) ^4–6^. According to the “slot” hypothesis, AMPARs associate with the PSD through the binding of their auxiliary subunit transmembrane AMPAR regulating proteins (TARPs) to PDZ-domain-containing scaffolding molecules in the PSD, including PSD-95 and other members of the membrane-associated guanylate kinase (MAGUK) family of proteins. As the PSD undergoes changes in organization and composition following the induction of synaptic plasticity, these PDZ domains can be dynamically occupied by AMPAR/TARP complexes and other transmembrane and non-transmembrane molecules ^6–10^.

One such non-transmembrane molecule is SynGAP, a synaptically-localized GTPase-activating protein (GAP) that negatively regulates small G-protein signaling important for activity-dependent changes in synaptic strength ^11–13^. SynGAP is an extremely abundant synaptic protein that is surpassed in copy number in the PSD by only the PSD-95 family of proteins and calcium/calmodulin-dependent protein kinase II (CaMKII) ^14^. Previously, we and others have shown that SynGAP undergoes a rapid and dynamic change in localization following neuronal activity ^7, 8^. At baseline, PSD-enriched SynGAP regulates synaptic plasticity by inhibiting several G-protein signaling cascades involved in LTP, including the Ras-Raf-MEK-ERK pathway, the activation of which is required for the insertion of AMPARs into the PSD ^15^. Following an LTP-inducing stimulus, SynGAP is phosphorylated by CaMKII in an NMDA-receptor-dependent manner and is rapidly dispersed from the PSD ^7^. SynGAP dispersion leads to increases in dendritic spine volume and synaptic AMPAR number ^7^. This dispersion relieves the negative regulation of synaptic Ras signaling and facilitates the induction and maintenance of physiological changes underlying activity-dependent synapse strengthening ^7^. Because SynGAP is highly abundant in the PSD, it is likely that SynGAP may occupy a significant number of the finite PDZ binding slots under basal conditions, which, in turn, may limit the number of “slots” for AMPAR/TARP complexes ^9^. Indeed, reduced SynGAP expression in heterozygous knockout (KO) mice have been reported to be associated with increased concentrations of TARPs and AMPARs within the PSDs of forebrain neurons in vivo ^9, 10^. However, *Syngap1* heterozygous mice also display enhanced activity of SynGAP- regulated downstream signaling pathways throughout development ^16^, making it difficult to know if the anti-correlation between SynGAP and TARP protein levels in the PSD is due to PDZ slot binding competition or changes in synaptic GAP activity and downstream signaling, or both.

Here, we report that SynGAP plays a key structural role at the PSD in regulating synaptic AMPAR number independent of its GAP activity. We tested this in vitro using catalytically inactive SynGAP expression constructs. We found that SynGAP GAP activity is not required for AMPAR recruitment to dendritic spines following LTP-dependent SynGAP dispersion in cultured neurons. To test the role of the GAP activity in vivo, we generated knock-in (KI) mice containing inactivating mutations within the SynGAP GAP domain. Surprisingly, hippocampal field recordings in brain slices from hetero and homozygote SynGAP GAP-activity-deficient KI mice (SynGAP^+/GAP*^ and SynGAP^GAP*/GAP*^) revealed that LTP expression was comparable to that observed in recordings obtained from their wild-type (WT) littermates. These findings contrast with those observed in heterozygous SynGAP KO mice (SynGAP^+/-^), which show significant LTP deficits relative to their WT littermates ^16, 17^. Moreover, the “GAP mutant” mice (SynGAP^+/GAP*^ and SynGAP^GAP*/GAP*^) also display normal activity, working memory, and contextual fear memory, in contrast to heterozygous SynGAP KO mice, which show significant impairments in these behaviors ^18, 19^. Further experiments demonstrate that SynGAP competes with TARP-γ8 for clustering in PSD-95-containing molecular condensates independent of its GAP activity and that cluster composition is regulated by SynGAP CaMKII phosphorylation sites. Taken together, these data provide strong evidence for a distinct structural role for SynGAP in the PSD that is independent of its role as a regulator of G-protein signaling and that SynGAP dispersion during LTP induction increases the number of PDZ-binding slots available for AMPAR/TARP complexes. These findings inform therapeutic strategies for the treatment of *SYNGAP1*-related neurodevelopmental disorders, as restoration of normal SynGAP downstream signaling pathways may not be sufficient to correct the aberrant AMPAR trafficking, plasticity, and circuit development associated with *SYNGAP1* haploinsufficiency.

## Results

### SynGAP GAP activity is not required for synaptic AMPAR recruitment in vitro

To test whether SynGAP regulates PSD composition in a GAP-independent manner, we employed a knockdown-replacement strategy in which we knockdown (KD) SynGAP with shRNA and replace it by transfecting wild-type (WT) or mutant SynGAP constructs in rat hippocampal neurons in vitro. We have previously used this approach to study SynGAP function during chemically induced LTP (cLTP) ^7^. cLTP causes SynGAP dispersion from the synapse, recruitment of AMPARs to synapses, and spine enlargement. In our previous study, we found that SynGAP KD increased synaptic Ras activity, enlarged spines, and increased levels of synaptic AMPARs in the basal state, which occluded further increases in spine size and receptor content upon cLTP induction^7^. Expression of WT SynGAP-α1 isoform rescued this phenotype. However, SynGAP harboring serine-to-alanine mutations at CaMKII phosphorylation sites critical for SynGAP dispersion from synapses (SynGAP 2SA; S1108A; S1138A) rescued the basal spine size and receptor content but failed to rescue cLTP due to deficits in SynGAP dispersion ^7^ (also see **Fig.1).** Here, we used this approach to examine the role of GAP activity in cLTP. We knocked down endogenous SynGAP by transfecting DIV19-21 rat hippocampal neurons with a short hairpin RNA against SynGAP (shRNA-SynGAP) and simultaneously expressing either a shRNA-resistant form of full-length Azurite-tagged WT or mutant SynGAP-α1 constructs, along with Super-ecliptic pHluorin (SEP)-tagged AMPAR GluA1 subunit (SEP-GluA1) and a mCherry cytosolic cell fill to observe SynGAP, AMPAR, and spine size changes during cLTP (**Fig. 1**). As observed previously, when SynGAP KD was rescued with shRNA-resistant WT SynGAP, cLTP stimulation resulted in the rapid dispersion of SynGAP from dendritic spines and a concomitant increase of both synaptic SEP-GluA1 signal and spine size (**Fig. 1*B, C, D***) ^7^. These cLTP-dependent changes were blocked when we transfected neurons with Azurite-SynGAP harboring the CaMKII phosphorylation site mutations. We then performed similar experiments using a SynGAP construct harboring two point mutations in the GAP domain at residues known to be critical for its GAP activity (SynGAP-GAP*; F484A, R485L) ^20, 21^. These mutations eliminated SynGAP GAP activity in a RAS activation pull-down assay in transfected HEK 293 cells (**Fig. S1**). SynGAP-GAP* was enriched at synapses and underwent dispersion from synapses following cLTP stimulation like WT SynGAP (**Fig. 1*B, C, D***). Remarkably, dispersion of SynGAP-GAP* was sufficient to rescue the cLTP-dependent enhancement of synaptic SEP-GluA1 signal, but not spine enlargement (**Fig. 1*B, C, D***). These data indicate that SynGAP regulates AMPAR synaptic accumulation during cLTP in a GAP-independent manner that is dissociable from the mechanisms underlying spine enlargement.

**Figure 1.**
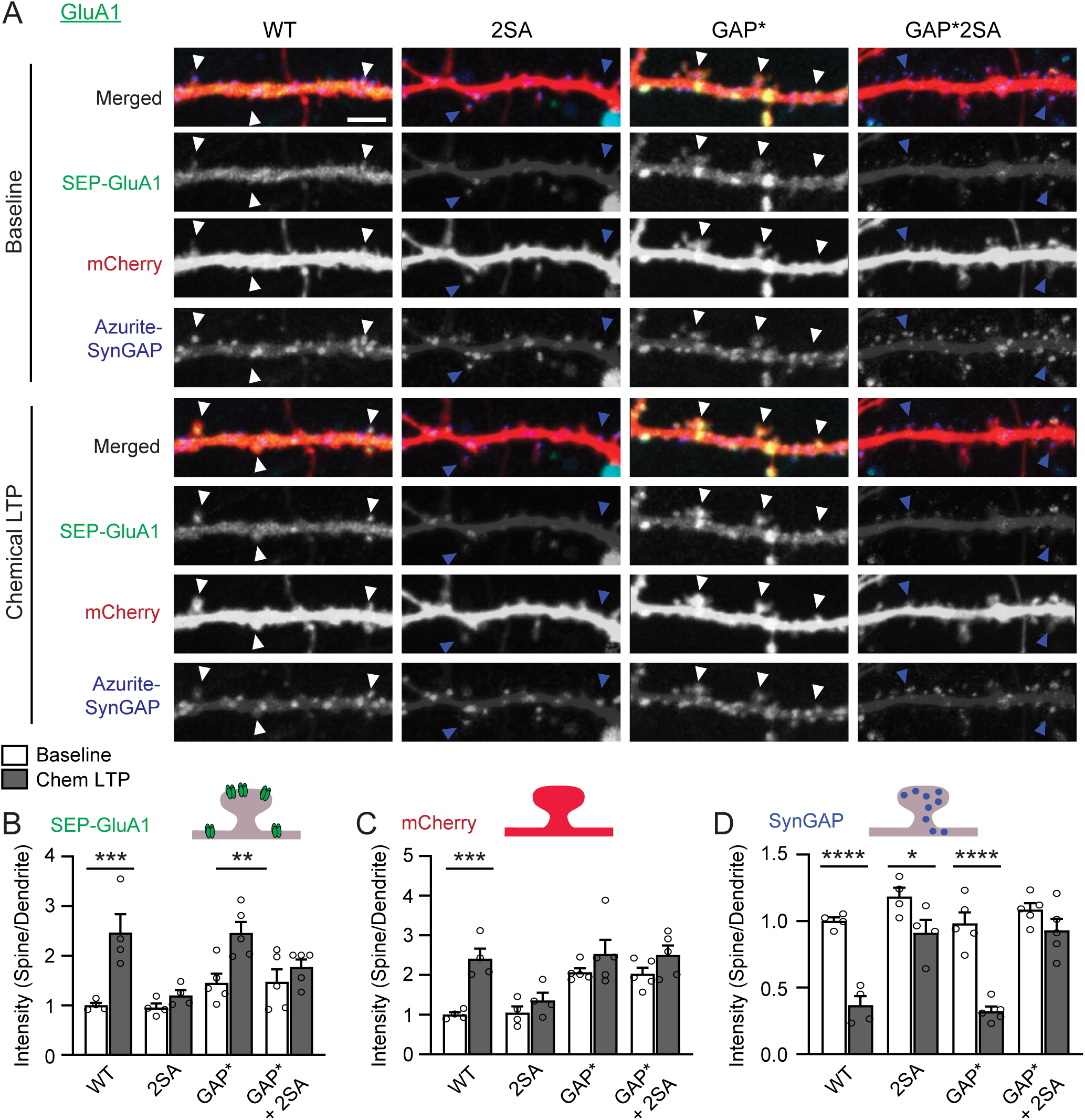
SynGAP GAP activity is not required for synaptic AMPAR recruitment in vitro. (A) Representative live fluorescent confocal images of a secondary dendrite from a rat hippocampal neuron transfected with mCherry (cytosolic cell fill), SEP-GluA1, and Azurite-tagged wild-type (WT) or mutant SynGAP before (Baseline) or after chemical LTP (cLTP). Mutants included phospho-deficient SynGAP (2SA), GAP-inactive SynGAP (GAP*), and a combination mutant with both (GAP*+2SA). Endogenous SynGAP was knocked down by shRNA and replaced by exogenous shRNA-resistant Azurite-SynGAP. Arrowheads indicate representative synaptic spine heads with SynGAP dispersion and SEP-GluA1 insertion. White arrowheads indicate dendritic spines that enlarge and exhibit SEP-GluA1 insertion and SynGAP dispersion in response to cLTP. Yellow arrowheads indicate dendritic spines displaying SEP-GluA1 insertion and SynGAP dispersion without spine enlargement. Blue arrowheads indicate spines with no response during cLTP. Scale Bar; 5 μm. (B) Quantification of SEP-GluA1 expression before and after cLTP in neurons transfected with WT and GAP* constructs. (WT: n = 4, Basal 1.000 ± 0.052 A.U., cLTP 2.465 ± 0.375 A.U.; 2SA: n = 4, Basal 0.956 ± 0.084 A.U., cLTP 1.200 ± 0.107 A.U.; GAP*: n = 5, Basal 1.453 ± 0.186 A.U., cLTP 2.459 ± 0.222 A.U.; GAP*+2SA: n = 5, Basal 1.472 ± 0.252 A.U., cLTP 1.772 ± 0.154 A.U.) (C) Quantification of the average change in spine volume during cLTP in neurons expressing WT and GAP* constructs, as measured by mCherry cell fill. (WT: n = 4, Basal 1.000 ± 0.059 A.U., cLTP 2.411 ± 0.253 A.U.; 2SA: n = 4, Basal 1.047 ± 0.163 A.U., cLTP 1.355 ± 0.199 A.U.; GAP*: n = 5, Basal 2.068 ± 0.100 A.U., cLTP 2.526 ± 0.359 A.U.; GAP*+2SA: n = 5, Basal 2.026 ± 0.161 A.U., cLTP 2.502 ± 0.242 A.U.) (D) Quantification of synaptic SynGAP expression before and after cLTP induction in neurons transfected with WT and GAP* constructs. (WT: n = 4, Basal 1.000 ± 0.028 A.U., cLTP 0.366 ± 0.070 A.U.; 2SA: n = 4, Basal 1.183 ± 0.067 A.U., cLTP 0.911 ± 0.099 A.U.; GAP*: n = 5, Basal 0.982 ± 0.083 A.U., cLTP 0.321 ± 0.037 A.U.; GAP*+2SA: n = 5, Basal 1.086 ± 0.049 A.U., cLTP 0.929 ± 0.088 A.U.) For Figure 1A-D: Two-way ANOVA with repeated measures for chemical LTP treatment, multiple comparisons with Sidak test. *p<0.05, **p<0.01, ***p<0.001, ****p<0.0001, n.s. (not significant).

### SynGAP-GAP-deficient mice have normal LTP

We next sought to determine whether the structural contribution of SynGAP to AMPAR trafficking we observed in vitro could be observed in vivo. To separate the role of G-protein signaling and the structural properties of SynGAP, we generated knock-in (KI) mice with the same inactivating mutations in the GAP domain used in vitro to eliminate GAP activity (**Fig. 2*A***). Heterozygote mice harboring this GAP* KI mutation (*Syngap1^+/GAP*^*) had normal levels of SynGAP protein expression in the brain, but showed increased levels of phosphorylated ERK (pERK) (**Fig. 2*B-G***), consistent with decreased GAP-activity. This has been shown previously in SynGAP heterozygous knock-out (KO) (*Syngap1*^+/-^) mice ^21^. Intriguingly, while homozygote SynGAP KO (*Syngap1^-/-^*) mice die perinatally within 2-3 days ^16, 17^, homozygote GAP* KI mice (*Syngap1^GAP*/GAP*^*) survive well beyond postnatal day 7 (**Fig. 2*H-I***), into adulthood, are fertile, and can be bred in homozygosity. Thus while SynGAP is required for viability, its GAP activity is not.

**Figure 2.**
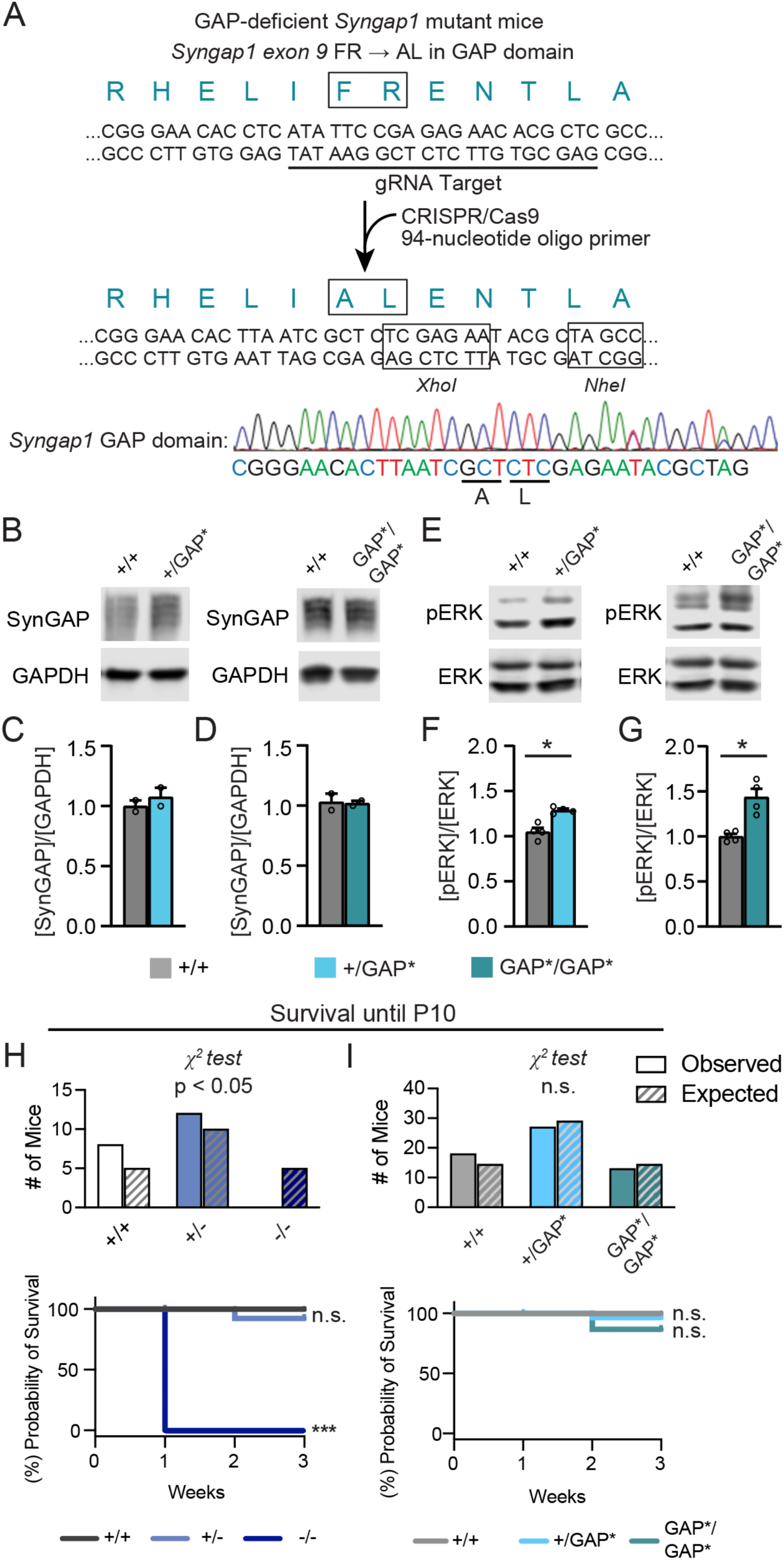
SynGAP-GAP-deficient KI mice exhibit normal SynGAP levels but have elevated Ras-ERK signaling in the brain. (A) Generation of *Syngap1^+/GAP*^* mice by CRISPR- Cas9. Guide RNA was designed to make the double-strand break near the target site, and the GAP activity-deficient mutant was introduced (FR to AL, “GAP*”) by homology-directed repair using a 94-nucleotide GAP-mutant oligo donor. (B, C, D) Representative immunoblots and quantification of SynGAP and GAPDH protein from whole brain of *Syngap1^+/GAP*^*, *Syngap1^GAP*/GAP*^*mice and wild-type littermates (*Syngap1^+/+^*). *Syngap1^+/+^*(n = 2, mean ± S.E.M.; 1.000 ± 0.046 A.U.) versus *Syngap1^+/GAP*^* (n = 2, mean ± S.E.M.; 1.076 ± 0.077 A.U.); *Syngap1^+/+^* (n = 2, mean ± S.E.M.; 1.030 ± 0.069 A.U.) versus *Syngap1^GAP*/GAP*^* (n = 2, mean ± S.E.M.; 1.020 ± 0.021 A.U.). Mann-Whitney test. p<0.05* (E, F, G) Representative immunoblots and quantification of phospho-ERK and total ERK protein from whole brain of *Syngap1^+/GAP*^*, *Syngap1^GAP*/GAP*^*mice and wild-type littermates (*Syngap1^+/+^*). *Syngap1^+/+^*(n = 4, mean ± S.E.M.; 1.05 ± 0.043 A.U.) versus *Syngap1^+/GAP*^*(n = 4, mean ± S.E.M.; 1.288 ± 0.017 A.U.); *Syngap1^+/+^* (n = 4, mean ± S.E.M.; 1.000 ± 0.029 A.U.) versus *Syngap1^GAP*/GAP*^* (n = 4, mean ± S.E.M.; 1.437 ± 0.092 A.U.). Mann-Whitney test. p<0.05* (H) Survival of *Syngap1^+/-^, Syngap1^-/-^* mice and wild-type littermates (*Syngap1^+/+^*) resultant from *Syngap1^+/-^*x *Syngap1^+/-^* breeding until age postnatal day 10 (P10). Top panel: Observed number of mice (*Syngap1^+/+^* = 8, *Syngap1^+/-^* = 12, *Syngap1*^-/-^ = 0) versus Expected number of mice (*Syngap1^+/+^* = 5, *Syngap1^+/-^* = 10, *Syngap1^-/-^* = 5); Chi-squared test, p<0.05*. No *Syngap1*^-/-^ mice survived until P10. Bottom panel: Survival plot. Log-rank (Mantel-cox) test; *Syngap1^+/+^*and *Syngap1^+/-^* (p = 0.36, n.s.); *Syngap1^+/+^*and *Syngap^-/-^* (p = 0.0009, ***). (I) Survival of *Syngap1^+/GAP*^*, *Syngap1^GAP*/GAP*^* mice and wild-type littermates c*Syngap1^+/+^*) resultant from *Syngap1^+/GAP*^*x *Syngap1^+/GAP*^* breeding until postnatal day 10 (P10). Top Panel: Observed number of mice (*Syngap1^+/+^* = 18, *Syngap1^+/GAP*^* = 27, *Syngap1^GAP*/GAP*^* = 13) versus Expected number of mice (*Syngap1^+/+^*= 14.5, *Syngap1^+/GAP*^* = 29, *Syngap1^GAP*/GAP*^* = 14.5; Chi-squared test, *n.s.* (p = 0.57). Bottom panel: Survival plot. Log-rank (Mantel-cox) test; *Syngap1^+/+^*and *Syngap1^+/GAP*^* (p = 0.41, n.s.); *Syngap1^+/+^*and *Syngap1^GAP*/GAP*^* (p = 0.11, n.s.).

To test whether SynGAP’s GAP activity is required for synaptic plasticity in vivo, we performed extracellular field recordings to measure long-term potentiation (LTP) in CA1 of the hippocampus induced by repeated theta-burst stimulation (TBS) of the Schaeffer collateral pathway in *Syngap1^+/-^*, *Syngap1^+/GAP*^*, and *Syngap1^GAP*/GAP*^*mice, along with their respective WT littermates (**Fig. 3**). We measured a 54% reduction of TBS- LTP expression in *Syngap1^+/-^*brain slices compared to WT littermates, replicating previously observed LTP deficits with *Syngap1* haploinsufficiency (**Fig. 3*A-B***) ^17^. In contrast, slices prepared from both *Syngap1^+/GAP*^*mice surprisingly exhibited normal TBS-LTP expression compared to recordings obtained from brain slices of WT littermates (**Fig. 3*C, D***). Even more remarkably, we found that *Syngap1^GAP*/GAP*^ mice* had normal LTP (**Fig. 3*C, D***). This data shows that the structural presence of SynGAP at synapses is sufficient for normal LTP to occur. Together, these data demonstrate that the GAP activity of SynGAP is not obligatory and is dispensable for the expression of hippocampal LTP.

**Figure 3.**
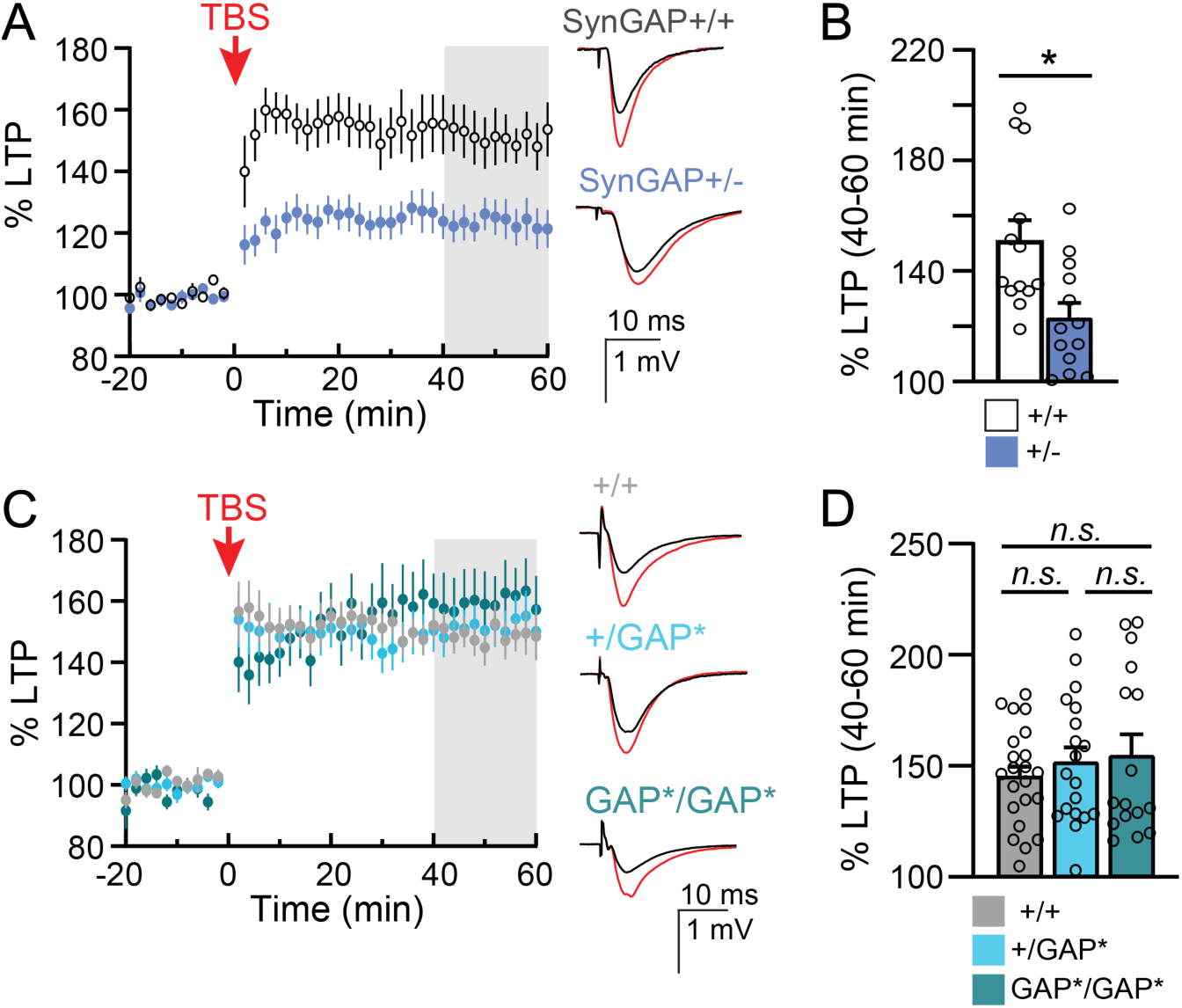
SynGAP-GAP-deficient mice have normal LTP. (A) Averaged population field CA1 recordings of TBS-LTP time course obtained from brain slices of *Syngap1^+/-^* mice and *Syngap1*^+/+^ littermate controls. All data points are normalized to the averaged baseline fEPSP slope. Inset: Example averaged fEPSP traces from *Syngap1*^+/+^ and *Syngap1^+/-^* slices recorded during baseline (black) and 40-60 minutes after TBS-LTP induction (red). (B) Quantification of averaged TBS-LTP in *Syngap1^+/-^* and *Syngap1*^+/+^ littermates Individual data points are superimposed. TBS-LTP is calculated by the ratio of the mean fEPSP slope measured 40-60 minutes after TBS-LTP induction (yellow-shaded region) divided by the averaged fEPSP baseline slope within each recorded sample. (*Syngap1*^+/+^: n = 13, 150.9 ± 7.51% S.E.M.; *Syngap1*^+/-^: n = 13, 123.0 ± 5.416% S.E.M.). Mann-Whitney ranked sum test. (C) Averaged population field CA1 recordings of TBS-LTP time course obtained from brain slices of *Syngap1*^+/GAP*^ and *Syngap1*^GAP*/GAP*^ mice as well as their *Syngap1*^WT-GAP^ littermate controls. Inset: Example averaged fEPSP traces from *Syngap1*^+/+^, *Syngap1*^+/GAP*^, and *Syngap1^GAP*/GAP*^* slices recorded during baseline (black) and 40-60 minutes after TBS-LTP induction (red). (D) Quantification of averaged TBS-LTP in *Syngap1*^+/+^*, Syngap1*^+/GAP*^, and *Syngap1*^GAP*/GAP*^ littermates. Individual data points are superimposed. (*Syngap1*^+/+^: n = 22, 145.5 ± 4.74% S.E.M.; *Syngap1*^+/GAP*^: n = 19, 151.9 ± 6.57% S.E.M.; *Syngap1^GAP*/GAP*^* : n = 16, 154.8 ± 9.34% S.E.M.). Non-parametric one-way ANOVA, Kruskal-Wallis multiple comparisons test. Error bars and shading represent the S.E.M.. p<0.05*, n.s. (not significant)

### SynGAP-GAP-deficient mice have normal activity, working memory, and associative fear memory

Previous work has shown that SynGAP is required for normal locomotor activity, learning, and memory ^18, 19, 22, 23^. However, whether the GAP activity of SynGAP is required for these behaviors has not been determined. To explore this, we performed a series of behavioral experiments in 2–4-month-old mice. In open field testing, *Syngap1*^+/-^ mice showed hyperactivity compared to WT littermates (**Fig. 4*A***), consistent with prior work ^22^.

**Figure 4.**
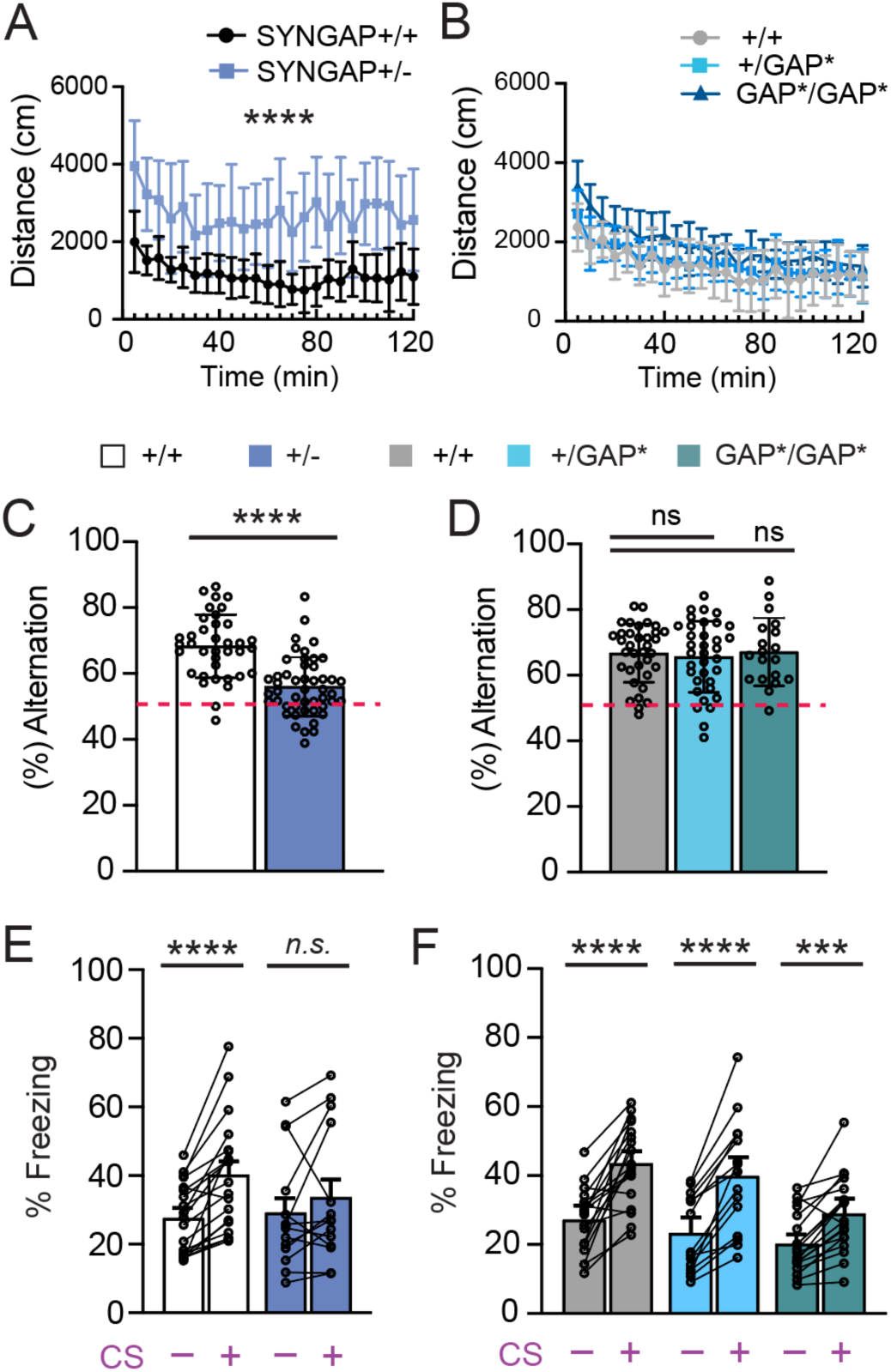
*Syngap1^+/GAP*^*KI mice have normal activity, working memory, and associative fear memory. (A) Distance traveled by *Syngap1^+/-^* mice (n = 15) and *Syngap1^+/+^* wild-type (WT) littermates (n = 18) during a 2-hour open field test in 5-minute intervals. Two-way ANOVA with repeated measures for time only, Šídák’s multiple comparisons test. (B) Distance traveled by *Syngap1^+/GAP*^* mice (n = 16), *Syngap1^GAP*/GAP*^* mice (n = 14), and *Syngap1^+/+^* WT littermates (n = 17) during a 2-hour open field test in 5-minute intervals. Two-way ANOVA with repeated measures for time only, Šídák’s multiple comparisons test. (C) Percentage of spontaneous alternating arm visits (% alternation) by *Syngap1^+/-^*mice (n = 48 56.00 ± 1.29% alternation) and *Syngap1^+/+^* littermates (n = 37, 68.30 ± 1.58% alternation) during a 5-minute Y-maze exploration test. The red dotted line represents the 50% successful alternation rate expected due to chance. Two-tailed student’s T-test. (D) Percentage of spontaneous alternating arm visits (% alternation) by *Syngap1^+/GAP*^*mice (n = 35, 65.63 ± 1.83% alternation), *Syngap1^GAP*/GAP*^* mice (n = 18, 67.14 ± 2.37 % alternation) and *Syngap1^+/+^* littermates (n = 35, 66.74 ± 1.49% alternation) during a 5-minute Y-maze exploration test. The red dotted line represents the 50% successful alternation rate expected due to chance. One-way ANOVA with Tukey test. (E) Average percentage of time spent freezing per minute (% freezing) with and without the conditioned stimulus (auditory cue, CS) by *Syngap1^+/-^* mice (n = 14 29.12 ± 4.44% freezing) and *Syngap1^+/+^* littermates (n = 19, 27.50 ± 2.37% freezing). Two-way ANOVA with repeated measures for CS only, Šídák’s multiple comparisons test. (F) Average percentage of time spent freezing per minute (% freezing) with and without presentation of the conditioned stimulus (auditory cue, CS) by *Syngap1^+/GAP*^*mice (n = 15, 23.18 ± 2.84% freezing), *Syngap1^GAP*/GAP*^* mice (n = 19, 20.11 ± 2.129 % freezing) and *Syngap1^+/+^* littermates (n = 18, 27.15 ± 2.203% freezing). Two-way ANOVA with repeated measures for CS only, Šídák’s multiple comparisons test. Error bars represent the S.E.M. *p<0.05, **p<0.01, ***p<0.001, ****p<0.0001, n.s. (not significant).

In contrast, both *Syngap1^+/GAP*^*and *Syngap1 ^GAP*/GAP*^* mice showed normal activity levels, indistinguishable from WT littermates (**Fig. 4*B***). We next compared working memory using the Y-maze spontaneous alternation task. Consistent with previous studies ^19, 23^, *Syngap1*^+/-^ mice had reduced spontaneous alternations compared to WT littermates (**Fig. 4*C***). However, the percent alternations for *Syngap1^+/GAP*^* and the *Syngap1 ^GAP*/GAP*^* mice were not significantly different from those of WT littermates (**Fig. 4*D***). To explore whether SynGAP GAP activity is required for associative learning, we then performed auditory cued and contextual fear conditioning. Consistent with prior studies ^18, 22^, *Syngap1*^+/-^ mice had impaired learning of a shock-associated auditory cue (conditioned stimulus; CS), as assessed by measuring the amount of time spent freezing in response to the presentation of the CS following conditioning (**Fig. 4*E***). Remarkably, *Syngap1^+/GAP*^* mice showed no significant impairment in fear conditioning and although *Syngap1 ^GAP*/GAP*^* mice had a trend of reduced conditioning compared to WT they still showed significant increases in freezing (**Fig. 4*F***). Taken together, these data show that while *Syngap1*^+/-^ mice exhibit hyperactivity and significant deficits in both working memory and fear learning, these impairments were not found in GAP-deficient heterozygous and homozygous GAP* KI mice.

### SynGAP competes with TARP-γ8 for binding to PSD-95 condensates

Since GAP catalytic activity of SynGAP is not required for normal LTP and memory, we wondered if SynGAP’s structural role in the PSD is essential for neuroplasticity. Both SynGAP ^24^ and TARP-γ8 ^25^ (hereafter referred to as γ8) undergo liquid-liquid phase separation (LLPS) with MAGUK family proteins in cell-free systems. LLPS is a known mechanism of the formation of molecular condensates that are comprised of dynamic protein clusters which exchange constituents with the adjacent pool of freely diffusing proteins ^26^. In vitro, many synaptic proteins are known to undergo LLPS, which can facilitate the clustering of membrane proteins ^27, 28^. Previous studies have shown that PSD-95 can form molecular condensates with both SynGAP ^24^ and TARPs^27, 28^. To explore whether this property of SynGAP is important for its role in LTP, we first investigated the possibility that SynGAP competes with γ8 for binding to PSD-95 to regulate the composition of synaptic PDS-95 molecular condensates during the expression of LTP. We transfected full-length PSD-95-mCherry and GFP-γ8 in HEK 293T cells in the absence and presence of SynGAP. Consistent with data from cell-free experiments ^25^, coexpression of PSD-95-mCherry and GFP-γ8 in the absence of SynGAP resulted in molecular condensates containing both GFP-γ8 and PSD-95-mCherry (**Fig. 5*A***). We then cotransfected GFP-γ8 and PSD-95-mCherry with increasing concentrations of SynGAP and examined the PSD-95 condensates for the presence of γ8 and SynGAP. At low levels of SynGAP1 expression, SynGAP co-clustered with γ8 and PSD-95, but with increasing concentrations of SynGAP, the presence of γ8 in the condensates was eliminated (**Fig. 5*A***), indicating that SynGAP can compete with γ8 for binding to PSD95 condensates.

**Figure 5.**
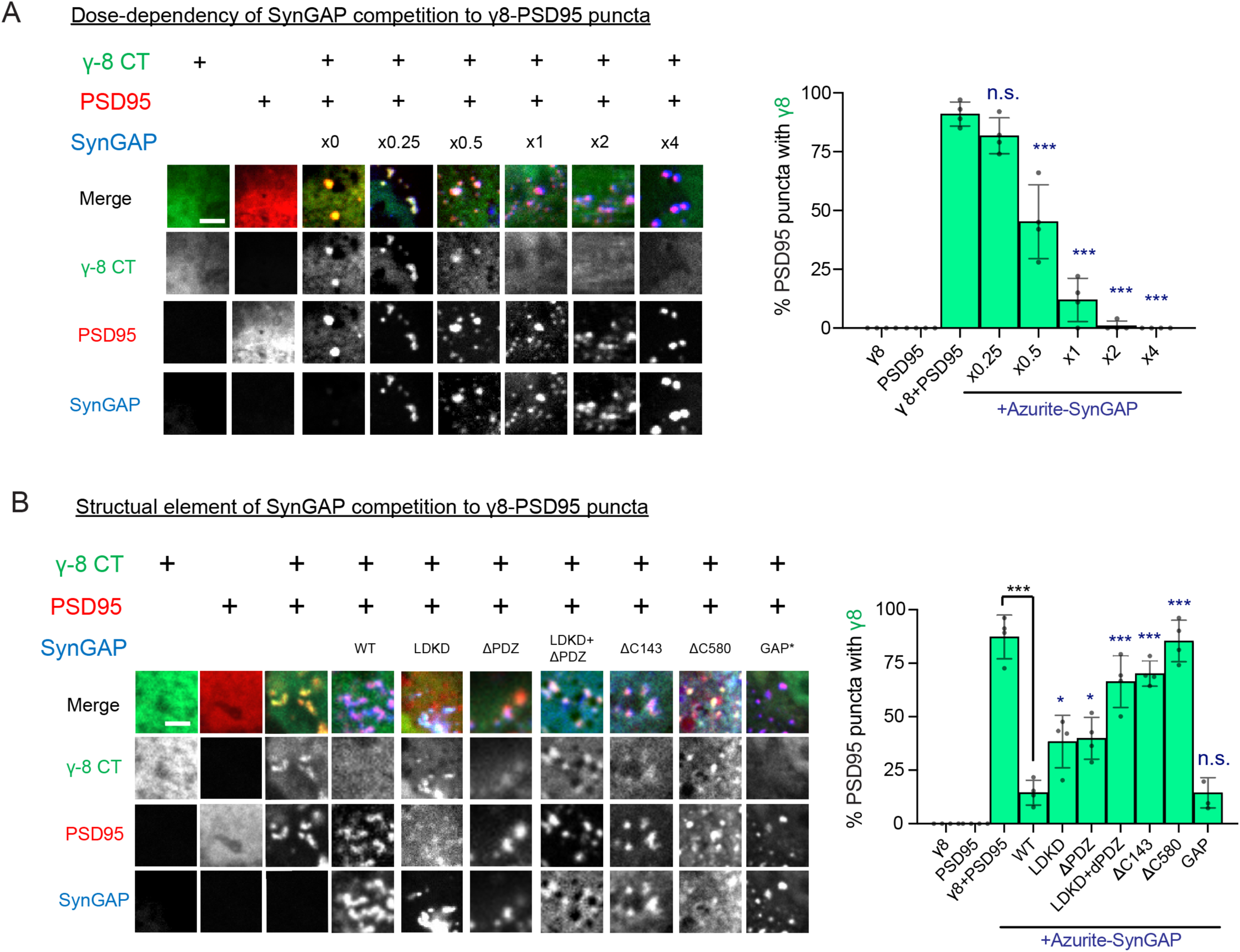
SynGAP-PSD95 and TARP-γ8-PSD95 compete in vitro. (A) Confocal microscopy of COS cells transfected with GFP-γ8, PSD95-mCherry, and different amounts of Azurite-SynGAP (x0.25, x0.5, x1, x2, x4). Scale bar; 5 μm. Right panel: (%) of PSD95 puncta with GFP-γ8 is displayed with different amounts of Azurite-SynGAP. One-way ANOVA with Tukey’s multiple comparisons test. Error bars represent the S.E.M. *p<0.05, **p<0.01, ***p<0.001, n.s. (not significant) compared to γ8+PSD95. (B) Confocal microscopy of COS cells transfected with GFP-γ8, PSD95-mCherry, and different Azurite-SynGAP mutants (WT, LDKD, ΔPDZ, LDKD+ΔPDZ, ΔC143, ΔC580, GAP*). Scale Bar; 5 μm. Right panel: (%) of PSD95 puncta with GFP- γ8 is displayed with different amounts of Azurite-SynGAP. One-way ANOVA with Tukey’s multiple comparisons test. Error bars represent the S.E.M. *p<0.05, **p<0.01, ***p<0.001, ****p<0.0001, n.s. (not significant) compared to γ8+PSD95+Azurite-SynGAP WT unless otherwise specified.

We then characterized the structural requirements of SynGAP1 for competition with γ8. Coexpression of WT SynGAP eliminated γ8 from PSD-95 condensates (**Fig. 5A**). While a series of mutations that regulate condensate formation had a significant impact on the ability of SynGAP to compete with γ8, a GAP-deficient mutation had no effect (**Fig. 5B**). Mutations in SynGAP’s PDZ ligand domain (ΔPDZ mutant) or in its coil-coil domain (LDKD mutant), which we have previously shown are important for condensate formation and LLPS ^24^, significantly decreased its ability to displace γ8 **(Fig. 5B)**. Combining these two mutations (1′PDZ/LDKD) almost completely eliminated SynGAP’s ability to compete with γ8. Deletion of the entire C-terminal coil-coil domain (1′C580) was required to eliminate SynGAP’s ability to displace γ8. These results indicate that SynGAP’s ability to undergo condensate formation and LLPS is essential for its ability to compete with γ8 for condensate formation with PSD-95. These results indicate that SynGAP’s c-terminal structure is essential for its ability to compete with γ8 for condensate formation with PSD- 95.

### Phase-in-phase separation of TARP-γ8-PSD95 and SynGAP-PSD95 condensates within phase-separated droplets of purified proteins

Next, we utilized purified proteins to further explore how γ8 and SynGAP compete for binding to PSD95. We first confirmed that γ8-PSD95 and SynGAP-PSD95 underwent LLPS and formed liquid condensates (droplets) in our assay system when the two pairs of purified proteins were mixed ^24, 25^ (**Fig. 6*A***; Top and Middle panels). Next, we explored the phase separation of these three purified proteins when combined. Interestingly, the proteins did not homogeneously mix within droplets and formed separate phase-in-phase condensates within droplets; SynGAP-PSD95 clustered in the center while γ8-PSD95 formed an adjacent protein condensation, a ring-like structure around the periphery (**Fig. 6*A***; Bottom panels). These distinct localizations were found in nearly all droplets (**Fig. S2*A***). When the localization of each protein is plotted on the DIC image, faint boundaries detected by refractive index changes were observed around the inner droplet of SynGAP- PSD95 in the DIC image (**Fig. 6*B***). This observation indicates that separate condensates with distinct light-scattering properties were forming within each droplet, with SynGAP- PSD95 condensates forming inside the γ8-PSD95 condensate. Plotting the localization of γ8-PSD95 by drawing a line across the droplet center when only γ8 and PSD-95 were mixed showed that γ8-PSD95 had a uniform distribution across the droplet (**Fig. 6*C* Left panel**). However, when SynGAP was added, γ8 was more peripherally localized with over 50% higher γ8 concentration in the periphery, while SynGAP was over 95% located centrally with PSD95 (**Fig. 6*C* Right panel**). This suggests that the γ8-PSD95 and SynGAP-PSD95 droplets form different layers of protein condensates.

**Figure 6.**
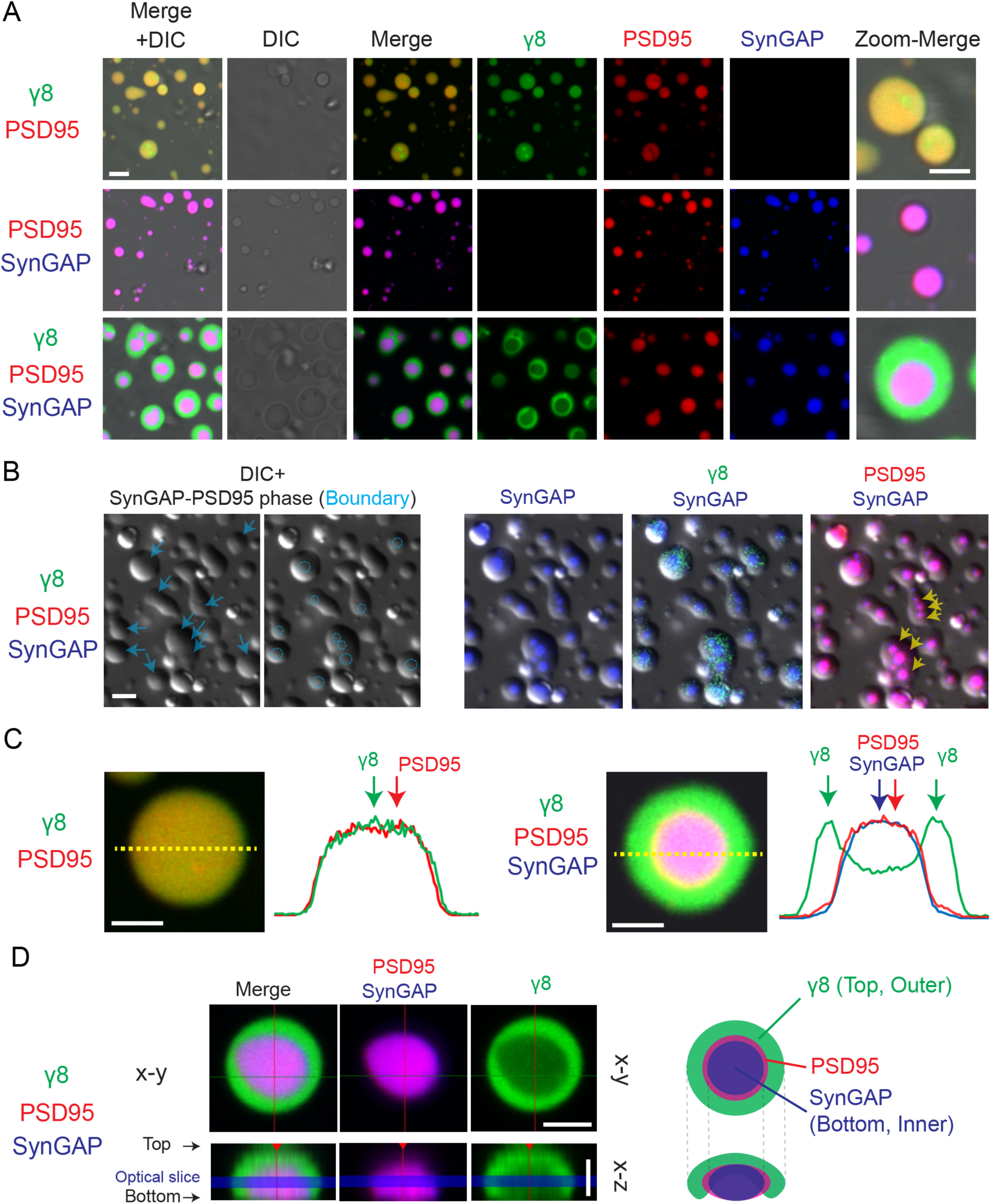
SynGAP-PSD95 and TARP-γ8-PSD95 show mutually exclusive phase-in-phase separation in droplets. (A) Images of purified protein sedimentation assay by confocal microscopy. Purified proteins included TARP-γ8 (“γ8”) tagged with iFlour568 (Green), PSD95 tagged with iFlour633 (Red), and SynGAP tagged with iFlour488 (Blue). Merge fluorescence images with Differential Interference Contrast (DIC) images (Left panels). γ8-PSD95 droplets (Top panels), SynGAP-PSD95 droplets (Middle panels), and γ8-PSD95-SynGAP droplets (Bottom panels). High-power views are also shown (right panels). Scale bar 3 μm. (B) Phase-in-phase separation of SynGAP-PSD95 droplets inside the γ8-PSD95 droplets. (Left panels) Blue arrows or circles delineate the inner rings of phase-in-phase separation. (Right panels) Merge of DIC images with γ8-PSD95-SynGAP droplets. Yellow arrows indicate regions of separation between SynGAP and PSD95 phase. Scale bar 3 μm. (C) Comparison between γ8-PSD95 droplets and γ8-PSD95-SynGAP droplets. A line scan of protein condensations (Yellow line) is shown to the right of each image. Scale bar 3 μm. (D) Optical sectioning microscopy of γ8-PSD95-SynGAP protein droplets. Top panels: x-y view. Bottom panels: x-z view. Optical slices (blue boxes) used to generate top (x-y) panels are shown. Scale bar 5 μm. Right panel: schematic of γ8- PSD95-SynGAP droplets.

Utilizing confocal microscopy to scan both the x-z and x-y planes, we found that the SynGAP-PSD95 droplet is not only located inside but also tends to sink closer to the bottom, likely due to its higher density. Conversely, γ8-PSD95 droplets are generally distributed outward and positioned in the plane above SynGAP-PSD95 (**Fig. 6*D***). Using time-lapse imaging, we investigated whether the droplets exhibit one of the fundamental properties of phase-separated bodies, the tendency to coalesce (**Fig. S2*B***). We observed that within 1-2 minutes after initial physical contact between droplets, the outer layer of γ8-PSD95 first fused, followed by the inner layer of SynGAP-PSD95. This observation suggests that both the outer γ8-PSD95 phase and the SynGAP-PSD95 phase retain their droplet-like properties.

This evidence strongly supports that SynGAP competes with γ8 for PSD95-binding resulting in the formation of the different layers of protein droplets rapidly and spontaneously. Critically, these phases appear incompatible with one another, exhibiting distinct droplet properties (e.g., density).

### Mutations in SynGAP that affect condensate formation and LLPS with PSD- 95 regulate recruitment of TARP-γ8 to synapses during cLTP

Next, we tested whether SynGAP GAP-activity is required for synaptic recruitment of γ8 during cLTP using the same SynGAP knockdown/replacement approach used above. Rat hippocampal neurons were transfected with GFP-γ8, and we observed a cLTP-dependent enhancement of synaptic GFP-γ8 fluorescence comparable to that observed with SEP-GluA1. Synaptic recruitment of γ8 required phosphorylation of SynGAP but did not require SynGAP GAP activity (**Fig. 7*A, B***), revealing that SynGAP regulates synaptic accumulation of γ8 during cLTP in a GAP-independent manner.

**Figure 7.**
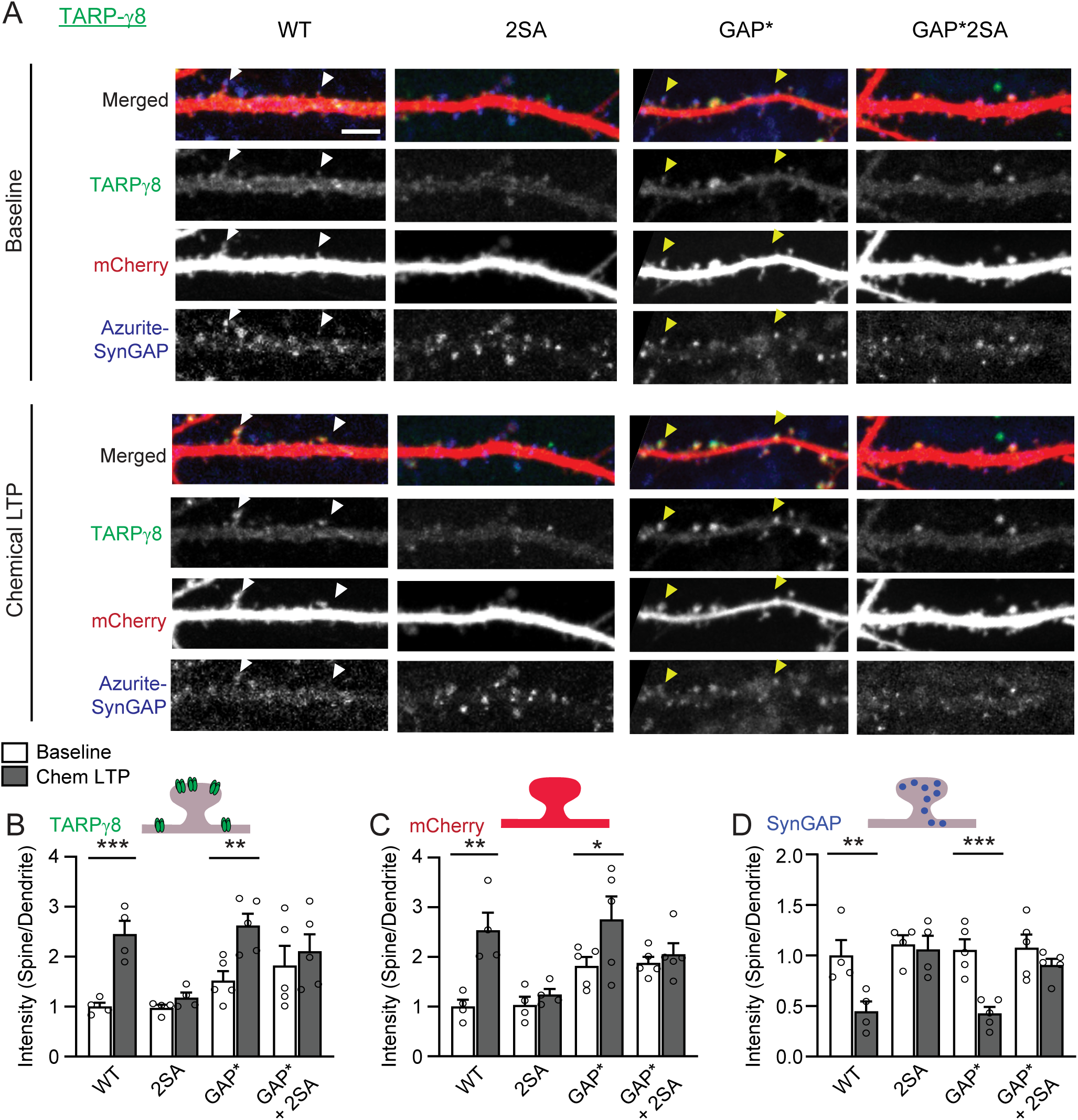
SynGAP GAP activity is not required for synaptic TARP-γ8 recruitment in vitro. (A) Representative live fluorescent confocal images of a secondary dendrite from a rat hippocampal neuron transfected with GFP-TARP-γ8, mCherry (cytosolic cell fill), and Azurite-tagged wild-type or mutant SynGAP before (Baseline) or after chemical LTP (cLTP). Mutants include phospho-deficient SynGAP (2SA), GAP-inactive SynGAP (GAP*), and a combination mutant with both (GAP*+2SA). Endogenous SynGAP was knocked down by shRNA and replaced by exogenous shRNA-resistant Azurite-SynGAP. Arrowheads indicate representative synaptic spine heads with SynGAP dispersion and γ8 insertion. White arrowheads indicate dendritic spines that enlarge and exhibit γ8 insertion and SynGAP dispersion in response to chemical LTP. Yellow arrowheads indicate dendritic spines displaying γ8 insertion and SynGAP dispersion without enlargement (no structural plasticity). Scale Bar 5 μm. (B) Quantification of synaptic GFP-γ8 expression before and after cLTP induction in neurons expression WT and GAP* constructs. (WT: n = 4, Basal 1.000 ± 0.079 A.U., cLTP 2.452 ± 0.265 A.U.; 2SA: n = 4, Basal 0.975 ± 0.064 A.U., cLTP 1.180 ± 0.103 A.U.; GAP*: n = 5, Basal 1.519 ± 0.190 A.U., cLTP 2.620 ± 0.238 A.U.; GAP*+2SA: n = 5, Basal 1.821 ± 0.393 A.U., cLTP 2.107 ± 0.342 A.U.) (C) Quantification of the average change in spine volume during cLTP in neurons expressing WT and GAP* constructs, as measured by mCherry cell fill. (WT: n = 4, Basal 1.000 ± 0.059 A.U., cLTP 2.411 ± 0.253 A.U.; 2SA: n = 4, Basal 1.047 ± 0.163 A.U., cLTP 1.355 ± 0.199 A.U.; GAP*: n = 5, Basal 2.068 ± 0.100 A.U., cLTP 2.526 ± 0.359 A.U.; GAP*+2SA: n = 5, Basal 2.026 ± 0.161 A.U., cLTP 2.502 ± 0.242 A.U.) (D) Quantification of synaptic SynGAP expression before and after cLTP induction in neurons transfected with WT and GAP* constructs. (WT: n = 4, Basal 1.000 ± 0.028 A.U., cLTP 0.366 ± 0.070 A.U.; 2SA: n = 4, Basal 1.183 ± 0.067 A.U., cLTP 0.911 ± 0.099 A.U.; GAP*: n = 5, Basal 0.982 ± 0.083 A.U., cLTP 0.321 ± 0.037 A.U.; GAP*+2SA: n = 5, Basal 1.086 ± 0.049 A.U., cLTP 0.929 ± 0.088 A.U.) Two-way ANOVA with repeated measures for chemical LTP treatment, multiple comparisons with Šídák’s test. *p<0.05, **p<0.01, ***p<0.001, ****p<0.0001, n.s. (not significant).

We then investigated if SynGAP mutations that alter SynGAP condensate formation with PSD-95 could affect the expression of cLTP. In these experiments, we knocked down SynGAP and replaced it with either WT SynGAP or SynGAP mutants that regulate LLPS. In these experiments, we induced cLTP at two glycine concentrations (10μM and 200μM) to test the sensitivity of SynGAP dispersion and cLTP induction to the strength of the induction stimulus. At 10μM glycine, WT SynGAP is not dispersed from spines, and cLTP was not expressed, as assayed by increases in spine size or the recruitment of γ8. In contrast, glycine at 200μM resulted in clear WT SynGAP dispersal and cLTP induction (**Fig. 8** and ^24^). Replacement with the LDKD mutant also rescued cLTP using 200μM glycine, while replacement with the PDZ mutant only partially rescued cLTP, highlighting the importance of PDZ ligand sequence of SynGAP for occupying PSD-95 PDZ domains in the basal state. Interestingly, at 10μM glycine, the LDKD mutant was dispersed and cLTP was expressed, in contrast to WT (**Fig. 8**). This result indicates that the LDKD mutation increases cLTP sensitivity to induction by glycine by decreasing the stimulation threshold for SynGAP dispersal from synapses and thus recruitment of γ8 during cLTP. Neither the 1′PDZ mutant nor the LDKD mutant exhibited significant effects on the synaptic targeting of SynGAP (**Fig. S3**). These results show that the LDKD mutation that modulates SynGAP’s ability to compete with γ8 for condensate formation with PSD-95 can regulate the threshold for recruitment of γ8 during cLTP induction, demonstrating that SynGAP’s ability to undergo LLPS is critical for LTP expression.

**Figure 8.**
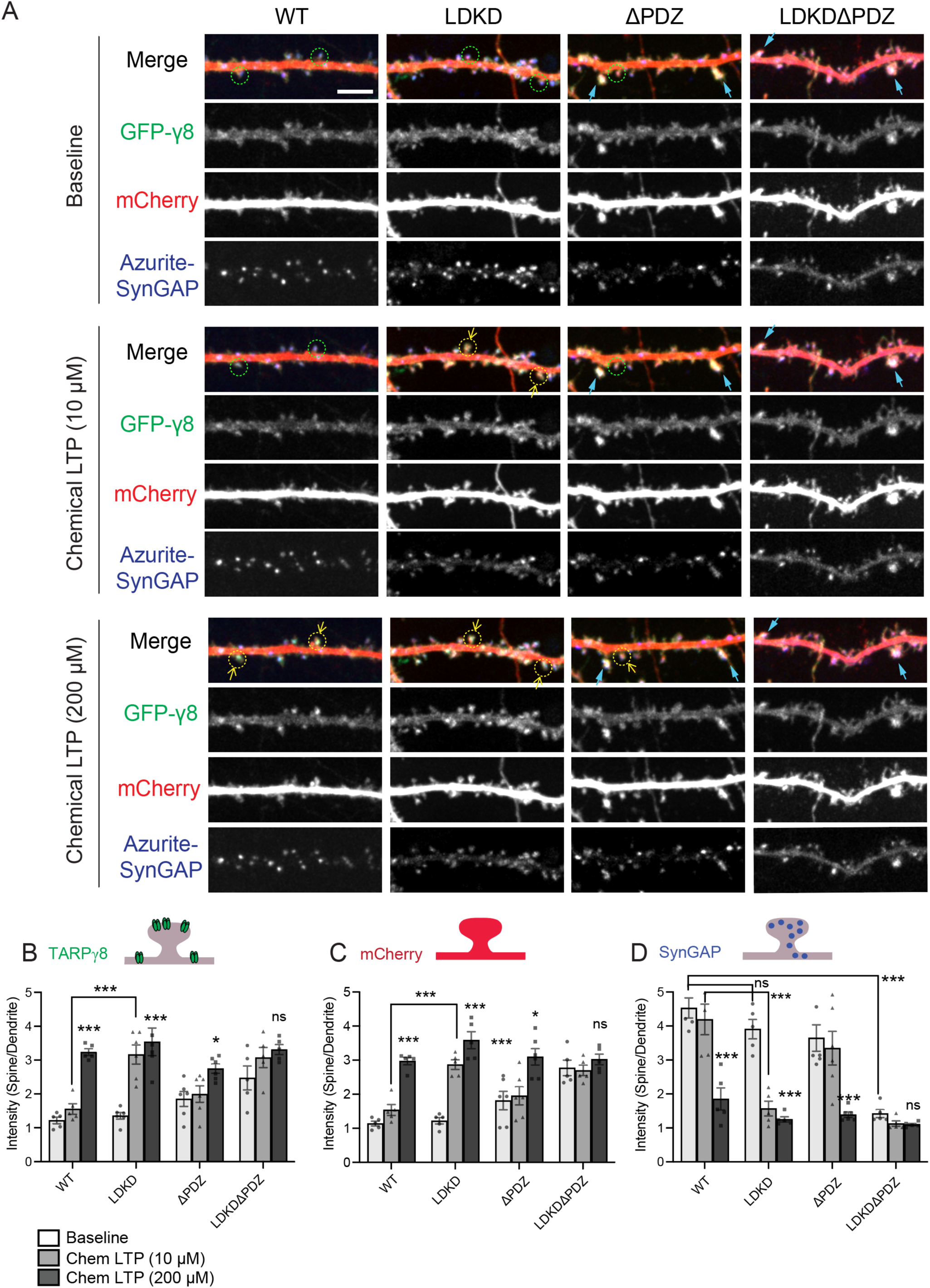
SynGAP phase-separation and PDZ-ligand binding capacity play crucial roles in the regulation of TARP- γ8 trafficking during LTP. (A) Representative live fluorescent confocal images of a secondary dendrite from a rat hippocampal neuron transfected with GFP-γ8, mCherry (cytosolic cell fill), and Azurite-tagged wild-type or mutant SynGAP before (Baseline) or after either weak cLTP (10 μΜ; Glycine) or strong cLTP (200 μΜ; Glycine). Mutants include LDKD, ΔPDZ, or both. Endogenous SynGAP was knocked down by shRNA and replaced by exogenous shRNA-resistant Azurite-SynGAP. Green circles indicate spine heads with the basal condition. Yellow circles and arrows indicate dendritic spines that enlarge and exhibit γ8 insertion, spine enlargements, and SynGAP dispersion in response to chemical LTP. Blue arrows indicate dendritic spines displaying γ8 insertion and large spine even in the basal state. Scale Bar; 5 μm. (B) Quantification of synaptic GFP-γ8 expression in secondary dendrites from rat hippocampal neurons transfected with WT, LDKD, ΔPDZ, and LDKD+ ΔPDZ constructs before and after cLTP. (WT: n = 5, Basal 1.218 ± 0.098 A.U. cLTP(10μΜ) 1.556 ± 0.157 A.U, cLTP(200μΜ) 3.237 ± 0.099 A.U.; LDKD: n = 6, Basal 1.356 ± 0.097 A.U. cLTP(10μΜ) 3.164 ± 0.285 A.U, cLTP(200μΜ) 3.540 ± 0.410 A.U. ΔPDZ: n = 6, Basal 1.853 ± 0.221 A.U., cLTP(10μΜ) 1.996 ± 0.243 A.U, cLTP(200μΜ) 2.748 ± 0.137 A.U.; LDKD+ΔPDZ: n = 5, Basal 2.474 ± 0.353 A.U., cLTP(10μΜ) 3.075 ± 0.300 A.U, cLTP(200μΜ) 3.312 ± 0.149 A.U.) (C) Quantification of the average change in spine volume as measured by mCherry cell fill in secondary dendrites from rat hippocampal neuron transfected with WT, LDKD, ΔPDZ, and LDKD+ ΔPDZ constructs before and after cLTP. (WT: n = 5, Basal 1.139 ± 0.070 A.U. cLTP(10μΜ) 1.538 ± 0.160 A.U, cLTP(200μΜ) 2.974 ± 0.109 A.U.; LDKD: n = 6, Basal 1.220 ± 0.092 A.U. cLTP(10μΜ) 2.868 ± 0.142 A.U, cLTP(200μΜ) 3.586 ± 0.249 A.U. ΔPDZ: n = 6, Basal 1.816 ± 0.270 A.U., cLTP(10μΜ) 1.954 ± 0.267 A.U, cLTP(200μΜ) 3.098 ± 0.242 A.U.; LDKD+ΔPDZ: n = 5, Basal 2.771 ± 0.225 A.U., cLTP(10μΜ) 2.698 ± 0.152 A.U, cLTP(200μΜ) 3.022 ± 0.156 A.U.) (D) Quantification of synaptic SynGAP expression in secondary dendrites from rat hippocampal neurons transfected with WT, LDKD, ΔPDZ, and LDKD+ ΔPDZ constructs before and after cLTP. (WT: n = 5, Basal 4.530 ± 0.296 A.U. cLTP(10μΜ) 4.194 ± 0.449 A.U, cLTP(200μΜ) 1.852 ± 0.326 A.U.; LDKD: n = 6, Basal 3.909 ± 0.284 A.U. cLTP(10μΜ) 1.569 ± 0.216 A.U, cLTP(200μΜ) 1.254 ± 0.075 A.U. ΔPDZ: n = 6, Basal 3.646 ± 0.389 A.U., cLTP(10μΜ) 3.348 ± 0.497 A.U, cLTP(200μΜ) 1.392 ± 0.080 A.U.; LDKD+ΔPDZ: n = 5, Basal 1.422 ± 0.120 A.U., cLTP(10μΜ) 1.128 ± 0.080 A.U, cLTP(200μΜ) 1.098 ± 0.033 A.U.) Two-way ANOVA with repeated measures for chemical LTP treatment, multiple comparisons with Šídák’s test. *p<0.05, **p<0.01, ***p<0.001, ****p<0.0001, n.s. (not significant).

## Discussion

The synaptic RasGAP SynGAP is essential for synaptic plasticity and learning and memory, and mutations in *SYNGAP1* cause intellectual disability, autistic-like behaviors, and epilepsy in humans. Recent studies have shown that the dispersion of SynGAP from synapses is required for the induction of LTP. We have previously demonstrated that SynGAP synaptic dispersion during cLTP relieves the basal inhibition of synaptic Ras, an important step that allows de-repression of ERK activity and AMPAR insertion ^7^. Whether SynGAP serves additional critical functions for AMPAR recruitment beyond its GAP activity has been an open question. Moreover, a comprehensive mechanistic understanding of how AMPARs are upregulated and maintained at the synapse during LTP has remained elusive. The slot hypothesis of LTP suggests that AMPAR/TARP complexes could bind to a finite number of available “slots” on scaffolding proteins at the PSD ^4–6, 9, 10^. Here, we provide evidence for the pivotal role of SynGAP in determining “slot” availability for the AMPAR/TARP complex independent of its GAP activity.

We had previously shown that phosphorylation of SynGAP by CaMKII is required for activity-dependent dispersion of SynGAP from the PSD during synaptic plasticity ^7^. Here, we reveal that AMPARs can be recruited to the PSD following SynGAP dispersion in a manner that is independent of the GAP activity of SynGAP. Additionally, we found that SynGAP binding to PSD-95 competes with TARPs for binding to PSD-95, and this antagonistic relationship is regulated by CaMKII phosphorylation sites on SynGAP. Taken together, these data indicate that CaMKII can act to molecularly tune PSD-95 binding partners at the PSD. Our data suggest that CaMKII may differentially regulate the affinities and condensation properties of SynGAP and TARPs for PSD-95 to promote the recruitment of AMPARs. The tuning of condensation properties is an attractive model to describe the rapid and dynamic changes to PSD composition and receptor density that occur during synaptic plasticity. Finally, we showed that the elimination of SynGAP GAP activity does not disrupt LTP in CA1 of the hippocampus, and several behavioral phenotypes are normal in *Syngap1* GAP mutant KI mice, suggesting that the structural function of SynGAP is a critical feature of its ability to regulate synaptic plasticity and to promote normal cognition.

This data demonstrate that SynGAP is a dominant driver of TARP enrichment in reconstituted condensates, as its CaMKII-dependent dispersion from synapses results in the recruitment of TARP-γ8 to synapses. It is known that CaMKII phosphorylates TARP C-terminal domains during synaptic plasticity ^6^. The phosphorylation of the TARP-γ2 C- terminus by CaMKII enhances its binding affinity for PSD-95 and AMPAR activity at synapses ^29^. Thus, it seems plausible that CaMKII phosphorylation of TARP C-termini contributes to our observed compositional switch phenomenon. However, we found that mutation of two key CaMKII sites on SynGAP eliminated the CaMKII-dependent dispersion of SynGAP and the recruitment of TARP-γ8. In addition, a recent report suggested that TARP-γ8 phosphorylation disrupts phase separation with PSD-95, resulting in decreased clustering ^25^. Additional experiments are required to explore the potential contribution of TARP phosphorylation in condensate composition-switching.

Previous studies have reported that the reduced SynGAP expression in heterozygous knockout KO mice is associated with increased concentrations of TARPs and AMPARs within the PSDs of forebrain neurons in vivo, suggesting a potential competition between SynGAP and TARPs for slots ^9, 10^. However, experimental results using *Syngap1 KO* heterozygous mice are confounded by the effects of persistent upregulation of synaptic small GTPase activity over the lifetimes of the animals tested. Here we have disentangled SynGAP signaling function from its structural properties both in vitro and in vivo by generating mice with inactivating GAP mutations. The heterozygous *Syngap1*^+/GAP*^ mice have reduced SynGAP GAP activity comparable to the heterozygous KO mice but have normal total SynGAP protein levels and displayed normal LTP and have no apparent deficits in several behaviors despite diminished GAP activity. Most remarkably, the homozygous *Syngap1*^GAP*/GAP*^ mice are viable and have normal LTP and behavior, indicating that LTP and viability are independent of the GAP activity. These results indicate that SynGAP-binding in the PSD is required for normal plasticity and cognition by regulating the number of PSD slots available for binding TARP/AMPAR complexes and, in turn, directly regulating synaptic strength.

These data do not suggest that the GAP-dependent signaling functions of SynGAP are unimportant. Further, work using these mice, and other approaches is needed to understand the role of SynGAP GAP activity in brain function.

## Conclusion

*SYNGAP1-*related ID has been classified as a RASopathy resulting from loss-of- function of the *SYNGAP1* gene ^30^. Several therapeutic strategies to ameliorate aberrant biochemical signaling downstream of Ras as a result of *SYNGAP1* haploinsufficiency have been tested ^31, 32^. However, the efficacy of this treatment approach remains inconclusive. Our new data suggest that pharmacologically correcting dysregulated downstream GAP signaling of SynGAP may not be sufficient to rescue disease phenotypes since these strategies do not address reduced PSD slot occupancy by SynGAP haploinsufficiency. Intriguingly, in searching surveys of various ages and ethnicities (a total of 687K entries encompassing GnomAD ^33^, TOPMED ^34^, 8.3KJPN ^35^ and ALFA), we found five human *SYNGAP1* single nucleotide variant carriers with GAP- disabling mutations ^20^ (rs1224277120 C>T: R485C, rs1248933822 G>A: R485H) that were not associated with any neurological or mental diagnosis. This suggests that heterozygous SynGAP mutations that impair GAP signaling are neither lethal nor sufficient to lead to an NDD diagnosis, unlike typical *SYNGAP1* loss-of-function mutations ^36, 37^ and is consistent with our results here. Our data indicate that future therapeutic strategies for the treatment of *SYNGAP1*-related ID should focus on increasing the amount of total SynGAP protein generated from the spared allele. These strategies will be complicated because SynGAP is expressed as a heterogeneous collection of structural isoforms that serve distinct functions in neuronal development and synaptic plasticity ^38^. Future studies will be needed to determine the structural and functional requirements for a complete rescue of *SYNGAP1* haploinsufficiency phenotypes, and this will help guide the development of treatments for *SYNGAP1-*related ID and other severe neurodevelopmental disorders.

## Acknowledgments

We thank all members of the Huganir lab for discussion and support throughout this work, especially Shaowen Ju, Ria Oba, Sam Myung, Sang Ho Kwon, and Stephanie Glavaris for reagent preparation and critical reading of the manuscript. We want to thank the Bridge the Gap SYNGAP Education and Research Foundation, the SynGAP Research Fund, and all the patients of SYNGAP1-related intellectual disability and their families for their outreach and advocacy.

## Author contributions

Conceptualization: YA, TG, RH

Methodology: YA, BL, IH

Investigation: YA, KR, EG, TG, RJ, THT

Formal Analysis: YA, KR, EG, TG, RJ, THT

Visualization: YA, KR, EG, TG

Funding acquisition: RH

Project administration: RH

Supervision: RH, AK

Writing – original draft: YA, KR, EG, TG, RH

Writing – review & editing: YA, EG, IH, RH

## Data and materials availability

All data are available in the main text or the supplementary materials.

## Supplementary Materials

Materials and Methods

Figs. S1 to S3

## Materials and Methods

### Molecular biology and cloning

All constructs used in this study were generated using standard restriction cloning protocols. GFP-TARP γ8 was generated by synthesizing a double-stranded gene fragment (gBlock™, IDT) with an engineered 5’ SalI site and a 3’ NotI site, followed by restriction subcloning into the CMV-driven GFP-C3 expression vector. For experiments involving purified proteins, coding sequences of proteins of interest were cloned into pGEX-6p bacterial expression vectors to generate protein products with an N-terminal GST tag with an engineered PreScission Protease recognition site for tag removal following isolation.

### Western blotting (SDS-PAGE)

5X SDS sample buffer (250 mM Tris, pH=6.8, 20% (v/v) Glycerol, 10% (w/v) SDS, 12% (v/v) b-Mercaptoethanol, 0.05% (w/v) Bromophenol Blue) was added to each sample for a final 1X concentration. Samples were sonicated with a probe sonicator before heating at 90°C for 5 minutes to facilitate complete protein denaturation. Samples were then loaded into pre-cast Bolt™ 4-12% gradient Bis-Tris, 1.0 mm gels (Invitrogen) soaked in commercially obtained Bolt™ MOPS or MES SDS running buffers (Invitrogen) depending on the molecular weights of proteins of interest. Proteins were separated on the basis of molecular weight by SDS PAGE. Proteins were then transferred to a 0.2 mm nitrocellulose membrane (Amersham Protran NC). Membranes were blocked with Intercept™ TBS blocking buffer (LI-COR Biosciences) for 30 minutes. Membranes were incubated with primary antibodies targeted to the proteins of interest in a solution of Tris-buffered saline with 0.1% Tween-20 (TBST) with 1.5% w/v bovine serum albumin (BSA) and 0.1% sodium azide overnight at 4°C with gentle rocking. Membranes were washed with TBST 3 times for 5 minutes before incubation with species-specific fluorescent secondary antibodies (LI-COR Biosciences) directed against bound primary antibodies for 1 hour at room temperature in the dark. Membranes were then washed with TBST 3 times for 10 minutes, followed by TBS 1 time for 5 minutes. Membranes were imaged using the Odyssey®CLx Imaging system. Protein bands were quantified using Image Studio software.

### Antibodies

GFP-tagged SynGAP, SynGAP_CT_, TARP-γ2, and TARP-γ8 were detected using a homemade primary antibody against GFP made in rabbits (JH4030) (1:1000 dilution). PSD-95 was detected using a monoclonal primary antibody against PSD-95 made in mice (isotype IgG2a) (NeuroMab) (1:2000 dilution). Phosphospecific SynGAP antibodies (pS1108, pS1138) produced in rabbits were used as described in ^7^ (1:1000 dilution). CaMKIIa was detected by a monoclonal antibody made in mice (6G9, ThermoFisher Scientific) (1:2000 dilution), and CaMKII pT286 was detected using a phospho-specific polyclonal antibody produced in rabbit (Abcam, ab5683) (1:2000 dilution). IRDye® 680RD or 800CW Donkey anti-Mouse IgG and Goat anti-Rabbit IgG secondary antibodies (LI-COR Biosciences) were used to detect primary antibodies of the appropriate species. Phospho-ERK and total-ERK were detected by the antibody obtained from Cell signaling (#9102 and #9101) (1:1000 dilution).

### Cell culture

HEK 293T cells originally obtained from ATCC (ATCC CRL-3216) were thawed from liquid nitrogen and maintained in a medium consisting of Dulbecco’s Modified Eagle Medium (DMEM) supplemented with 10% Fetal Bovine Serum (FBS) (Hyclone), and 1% Penicillin-Streptomycin (10,000 U/mL) (Thermo Fisher Scientific). Cells were maintained at 37°C in an incubator with 5% CO_2_. Cells were passaged fewer than 20 total times.

Cultured primary hippocampal neurons were prepared as described previously ^7^, with some modifications. Briefly, hippocampi were dissected from embryonic day 18 (E18) rat pups before dissociation by papain treatment and mechanical trituration. Cells were plated on coverslips treated with 25 mM poly-L-Lysine in Neurobasal media (Gibco) supplemented with 5% horse serum (Hyclone), 1% Penicillin-Streptomycin (10,000 U/mL) (Thermo Fisher Scientific), 2 mM GlutaMAX Supplement (Thermo Fisher), 2% B27 supplement (Gibco) (NM5). One day following plating, the medium was replaced with medium lacking horse serum (NM0). Neurons were maintained in an incubator at 37°C with 5% CO_2_ for up to 21 days, and media (NM0) was changed once per week over the duration of the culture.

### Confocal live cell imaging

Live HEK 293T cells and cultured hippocampal neurons were imaged using a Cell Observer spinning disk confocal microscope (Carl Zeiss) or an LSM 880 laser scanning confocal microscope (Carl Zeiss). For live imaging of HEK 293T cells, cells were plated on collagen-coated 18 mm glass coverslips 24-36 hours before the start of the experiment. HEK 293T cells were transiently transfected with cDNA constructs encoding proteins of interest using Lipofectamine 2000 (Invitrogen) 16-24 hours before the start of the experiment. Coverslips containing cells were pre-incubated for 20 minutes in basal extracellular solution (ECS) (150 mM NaCl, 3 mM KCl, 2 mM CaCl_2_, 10 mM HEPES pH=7.4, 10 mM D-(+)-Glucose, 1 mM MgCl_2_) at 37°C with 0% CO_2_ before being transferred to a custom-made imaging chamber filled with basal ECS. Experiments were performed at 37°C. Cultured hippocampal neurons were transiently transfected with the desired cDNA constructs using Lipofectamine 2000 at DIV17-19, 36-48 hours preceding the start of the experiment. Coverslips containing cultured neurons were subjected to the same pre-incubation protocol applied to HEK 293T cells before being loaded into the imaging chamber.

### Protein purification

GST-tagged SynGAP CC-PBM, PSD95 full length^24^, γ8 C-terminal tail^25^ were expressed in Escherichia coli BL21 in LB medium at 37°C for around 3 hours. Protein expression was induced by 0.1 mM IPTG (final concentration) at OD_600_ around 1.0 at 30°C. Proteins were purified using glutathione Agarose affinity column, and GST tags were cleaved by PreScission Protease (Cytiva 27-0843-01) with a cleavage buffer containing 50 mM Hepes pH 7.4, 50 mM NaCl, 1 mM EDTA, 1 mM DTT. Purified proteins eluted from the affinity column were then collected and measured the concentrations by Nanodrop. γ8 proteins were further concentrated using Amicon Ultra centrifugal filter with MWCO 3K (Merck Millipore UFC500324).

### Protein fluorescence labeling and an imaging-based assay of phase separation

Purified proteins were labeled by iFluor-488/568/633 succinimidyl ester (AAT Bioquest 1023/1049/1030) as previously described ^28^. Fluorescence-labeled protein was mixed with corresponding unlabeled protein at 1:20. For imaging assay, γ8 C-terminal tail (50 μM) and PSD95 full-length (10 μM), and SynGAP CC-PBM (10 μM) were mixed in cleavage buffer plus 5% PEG and total volume of 10 μl. Each condition mixture was injected into a flowmetry chamber (comprised of a #1.5H coverslip (Carl Zeiss 474030- 9000) and a slide glass separated by two layers of double-sided tape as a spacer). Zeiss LSM880 confocal microscope (20X Plan-Apochromat M27, Air, NA 0.8, or 63X Plan-Apochromat M27, Oil, NA 1.4, Carl Zeiss) was used for DIC and fluorescent imaging at room temperature. The ImageJ software was used for analyzing images.

### Chemical LTP (cLTP) stimulation

Live imaging and quantification of LTP were performed as described previously ^7^ Hippocampal neurons from embryonic day 18 (E18) rats were seeded on 25-mm poly-L-lysine-coated coverslips. The cells were plated in Neurobasal media (Gibco) containing 50U/ml penicillin, 50mg/ml streptomycin, and 2 mM GlutaMax supplemented with 2% B27 (Gibco) and 5% horse serum (Hyclone). At DIV 6, cells were thereafter maintained in glia-conditioned NM1 (Neurobasal media with 2mM GlutaMax, 1% FBS, 2% B27, 1 x FDU (5mM Uridine (SIGMA F0503), 5 mM 5-Fluro-2’-deoxyuridine (SIGMA U3003). Cells were transfected at DIV17-19 with Lipofectamine 2000 (Invitrogen) in accordance with the manufacturer’s manual. After two days, coverslips were placed on a custom perfusion chamber with basal ECS (143 mM NaCl, 5 mM KCl, 10 mM Hepes pH 7.42, 10 mM Glucose, 2 mM CaCl_2_, 1 mM MgCl_2_, 0.5 μM TTX, 1 μM Strychnine, 20 μM Bicuculline), and time-lapse images were acquired with either LSM880 (Carl Zeiss) or Spinning disk confocal microscopes controlled by Axiovision software (Carl Zeiss). Following 5-10 min of baseline recording, cells were perfused with 10 ml of glycine/ 0Mg ECS (143 mM NaCl, 5 mM KCl, 10 mM HEPES pH 7.42, 10 mM Glucose, 2 mM CaCl_2_, 0 mM MgCl_2_, 0.5 μM TTX, 1 μM Strychnine, 20 μM Bicuculline, 200 μM Glycine) for 10 min, followed by 10 ml of basal ECS. To stabilize the imaging focal plane for long-term experiments, we employed Definite focus (Zeiss). For quantification, we selected pyramidal neurons based on morphology that consisted of a clear primary dendrite and quantified all spines on the 30-40 μm stretch of the secondary dendrite beginning just after the branch from the primary dendrite. For identifying spine regions, we used the mCherry channel to select the spine region that was well separated from dendritic shaft. These regions of interest (ROIs) in the mCherry channel were transferred to the green channel to quantify total SynGAP content in spines. Total spine volume was calculated as follows; (Average Red signal at ROI – Average Red signal at Background region) * (Area of ROI). Total SynGAP content was calculated as follows; (Average Green signal at ROI – Average Green signal at Background region) * (Area of ROI). Through this quantification, we can precisely quantify the total signals at each spine, even if the circled region contained some background area. For [%] spine enlargement before/ after LTP, we took a relative ratio of the total spine volume (total red signal) of each spine before/ after LTP ([%] spine enlargement = (Total Red Signal after cLTP / Total Red signal at basal state-1)*100). For [%] SynGAP dispersion, we calculated the degree of total SynGAP content loss after cLTP at each spine compared to the total SynGAP content at basal state ([%] dispersion = (1- Total Green Signal after CLTP / Total Green signal at basal state) * 100).

### Generation of Syngap1-GAP-deficient knock-in mice

All animals (rats and mice) used in this study were housed in the JHU SOM animal facility according to JHU Institutional Animal Care and Use Committee (IACUC) guidelines. SynGAP^+/-^ knockout mice were generated using a BAC transgene at the Transgenic Core Facility of JHU SOM ^17^. Mice were backcrossed onto the C57BL/6 background. SynGAP^+/GAP*^ and SynGAP^GAP*/GAP*^ mice were engineered using CRISPR/Cas9 genome editing resulting in two amino acid changes (F484A, R485L) on a mixed C57BL/6J background. Cas9 was targeted to the genomic sequence of interest using the following guide RNA (gRNA) sequence: 5’-GCGTGTTCTCTCGGAATATG-3’. The following 94 bp edited oligo donor template containing 9-point mutations (5 silent, 4 to generate FR→AL mutations) was introduced: 5’-ATCAGTCTCATATACTCTTCTATGGCTTTAGTGGCTAGCGTATTCTCGAG AGCGATTAAGTGTTCCCGCTCCATGAATCGGTCTACCTCTGACA-3’.

The protospacer adjacent motif (PAM) was destroyed, and two restriction sites (XhoI and NheI) were engineered downstream of the FR→AL mutations for founder screening.

### Animals

SynGAP heterozygous knockout mice (previously described Kim et al., 2003), SynGAP^GAP*^ mutants, and wild-type (WT, WT-GAP) littermates were maintained on a mixed background of C57/B6J and 129/SvEv background strains. Animals were allowed *ad libitum* access to food and water and reared on a typical 12-hour light-dark cycle. All animal experiments utilized both male and female mice (aged 3-7 months) and were conducted in accordance with the guidelines implemented by the Institutional Animal Care and Use Committee at Johns Hopkins University.

### Acute slice preparation

*SynGAP^+/GAP*^, SynGAP^GAP*/GAP*^,* and *SynGAP^+/-^* mice (3-7 months of age), along with their respective wild-type (WT-GAP, *Syngap1^+/+^*) littermates, were transcardially perfused with ice-cold aerated dissection buffer (212.7 mM sucrose, 5 mM KCl, 1.25 mM Na_2_PO_4_, 10 mM glucose, 26 mM NaHCO_3_, 0.5 mM CaCl_2_, 10 mM MgCl_2_) under isoflurane anesthesia immediately before decapitation. The brain was rapidly removed from the skull, and the hippocampi were extracted in a continuously oxygenated dissection buffer. Acute transverse hippocampal slices (400 µm thickness) were prepared using a vibratome (Leica VT1200S) and briefly washed of the sucrose-based dissection buffer in aerated artificial cerebrospinal fluid (ACSF) composed of (in mM): 119 NaCl, 5 KCl, 1.25 Na_2_PO_4_, 26 NaHCO_3_, 10 glucose, 2.5 CaCl_2_, and 1.5 MgCl_2_. Slices were then placed in a chamber containing ACSF at 30°C for 30 minutes, and then transferred to room temperature for at least 60 minutes until used for electrophysiological recordings. The experimenter was blinded to the animals’ genotypes until all experiments and analyses were completed.

### Extracellular LTP recordings

Slices were placed in a submersion recording chamber with recirculating aerated ACSF at 30°C. Synaptic field excitatory postsynaptic potentials (fEPSPs) were evoked in response to electrical stimulation of the Schaeffer collateral inputs via a bipolar theta glass Ag/AgCl electrode (3 MΩ) containing ACSF. Input-output curves were used to determine the half-maximal fEPSP amplitude, which was the stimulation intensity used to measure the fEPSP slope over a stable 20-minute baseline period in response to a single 0.2-ms stimulation pulse delivered every 30 seconds. (A stable baseline period of 10 minutes with baseline fEPSP slope not drifting by >10% was the minimum required inclusion criteria for LTP recordings.) To induce LTP, four episodes of theta-burst stimulation (TBS) were triggered at 0.1 Hz. Each TBS episode consisted of 10 stimulus trains administered at 5 Hz, whereby one train consisted of 4 pulses at 100 Hz. Following TBS, the fEPSP slope was measured for 60 minutes by delivering single electrical pulses every 30 seconds. The magnitude of LTP was quantified by normalizing the fEPSP slope to the average baseline response, then calculating the average fEPSP slope between 40-60 min after TBS. Statistical comparisons were made exclusively between WT and mutant littermates with a student T-test, Mann-Whitney test, or one-way ANOVA.

### Behavior

Mice aged 2-4 months were subjected to behavioral tests, including Open Field to assess locomotion, the Y-maze spontaneous alternation task to assess working memory performance, and Contextual Plus Cued Fear Conditioning to assess associative memory. All groups were approximately evenly divided (45-55%) between males and females. All animals were housed in the JHU SOM Miller Research Building (MRB) animal facility. All behavioral assays were performed in the JHU School of Medicine Animal Behavioral Core.

### Open field task

Each test mouse was placed in a photobeam–equipped (16 x 16 configuration with equal spacing of 2.54 cm) plastic chamber (45 × 45 cm) and was allowed to explore free from interference for 120 minutes. Ambulatory movements were tracked and analyzed using the Photobeam Activity System – Open Field (San Diego Instruments).

### Y-maze spontaneous alternation task

Following a 30-minute acclimatization period, mice were placed in the center of a three-chamber Y-maze in which the three arms were oriented 120 degrees from one another. Mice were allowed to explore the apparatus for 5 minutes. Arm entries were recorded when both of the mouse’s rear paws passed over the boundary line between the apparatus’s center region and arm region. An arm entry was recorded as an alternation when the mouse fully entered an arm that it had not visited most recently (e.g., Arm A → Arm B → Arm C = Alternation; Arm A → Arm B → Arm A = Not Alternation). % Alternation was calculated as the number of alternating arm entries divided by the total number of arm entries. The Y-maze apparatus was thoroughly cleaned between trials. Arm entries were recorded manually, and the experimenter was blind to the experimental conditions. The normality of the data was assessed using the Kolmogorov-Smirnov test, and groups were statistically compared using a one-way ANOVA followed by Tukey’s post hoc multiple comparisons test.

### Contextual and Cued Fear Conditioning

Contextual and cued fear conditioning test was performed as previously described ^18^. Animals were handled (picked up and held by the experimenter for 30s) daily for 14 days before training. Before this test, mice were kept in a soundproof room separate from the testing room for 30 minutes. To assess fear-related learning and memory, mice were placed singly in an acrylic chamber of PVC plastic walls (33 × 25 × 28 cm) with a stainless-steel grid floor (0.2 cm diameter, spaced 0.5 cm apart). Between mice for conditioning and context testing, the walls and grids of the chamber were wiped with 70% ethanol. In the cued test, the walls and floor were cleaned with 1% acetic acid. Animals were placed in the conditioning chamber (Video Freeze; Med Associates) and allowed to explore for 120 seconds, after which a 30 s white noise tone (90 dB) was presented, which co-terminated with a footshock (1s, 0.5 mA). This tone–shock pairing occurred three times per session, with an intertrial interval of 90 seconds. After the third and final shock, the animals remained in the training chamber for 90 seconds. Twenty-four hours later, the animals were returned to the chamber. Contextual memory was assessed by the percent of time spent freezing for 600 seconds (no shock presentation). Twenty-four hours later, the dimensions, as well as the visual, tactile, and olfactory cues (cleaning with 1% acetic acid) of the conditioning chamber, were altered to create a novel context for the mice. Cued fear learning was assessed in this novel context by measuring the average % freezing per minute for 300s without and 300s with the presentation of the auditory cue (conditioned stimulus, CS).

### Statistics

All statistical analyses were performed using GraphPad Prism 8 software. Graphs were prepared using GraphPad Prism 8.0 Software or Microsoft Excel 16.36. All error bars and shadows represent the standard error of the mean (S.E.M.) unless otherwise stated. The normality of the data was assessed using the Shapiro-Wilk test unless stated otherwise. Western blot data were analyzed using a one-way ANOVA followed by Tukey’s posthoc multiple comparisons test unless stated otherwise. FRAP data were analyzed by nonlinear regression and were fit to one-phase functions. When applicable, curve plateaus were compared using the extra sum-of-squares F test. Data obtained from the ionophore stimulation experiments were analyzed using a two-way RM-ANOVA followed by Šídák’s posthoc multiple comparisons test unless otherwise stated. For all behavior testing, the normality of the data was assessed using the Kolmogorov-Smirnov test. In the Y-maze spontaneous alternation task, groups were compared using a one-way ANOVA followed by Tukey’s post hoc multiple comparisons test. For Context-Cue Fear-conditioning, mutant mice were compared to their WT littermates during the Context trial using a one-way repeated measures ANOVA. With and without cue was compared within each genotype using a one-way ANOVA. *p<0.05, **p<0.01, ***p<0.001, ****p<0.0001.^15^

---

**Figure S1.**
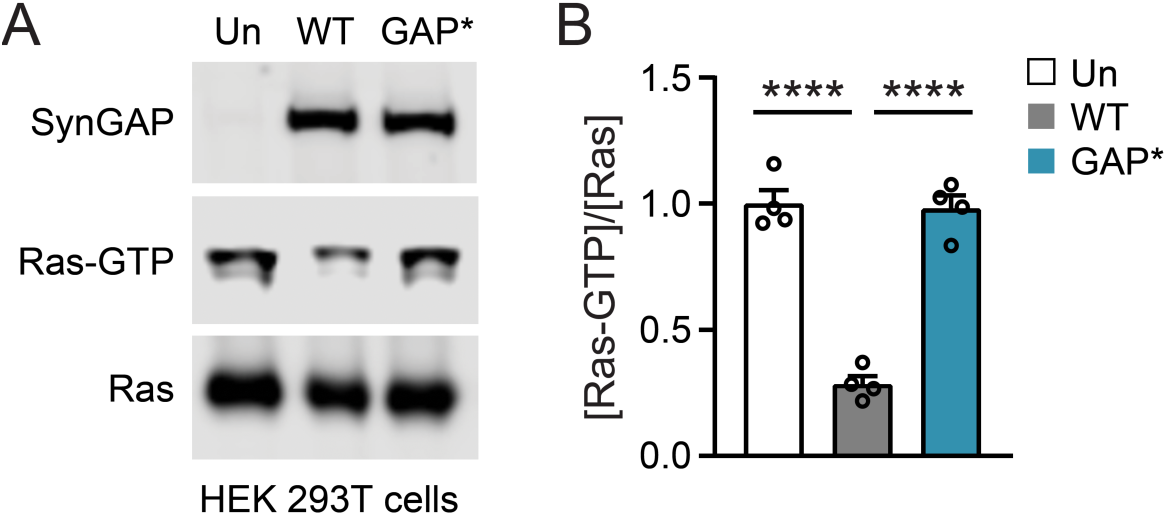
Expression of GAP-deficient SynGAP in HEK 293T cells lacks GAP activity and does not alter SynGAP expression. (A) Representative Western blots of SynGAP, Ras-GTP, and Ras in whole-cell lysates prepared from untransfected HEK 293T cells or HEK 293T cells that overexpress SynGAP-WT or SynGAP-GAP*. (B) Quantification of activated Ras (Ras-GTP/Ras) in the Western blot shown in S1A. (Untransfected: n = 4, 1.000 ± 0.054 A.U.; SynGAP-WT: n = 4, 0.285 ± 0.0319 A.U.; SynGAP-GAP*: n = 4, 0.981 ± 0.052 A.U.) Statistics: One-way ANOVA with Tukey’s multiple comparisons test. p<0.05*, < 0.01**, <0.001***, < 0.0001****

**Figure S2.**
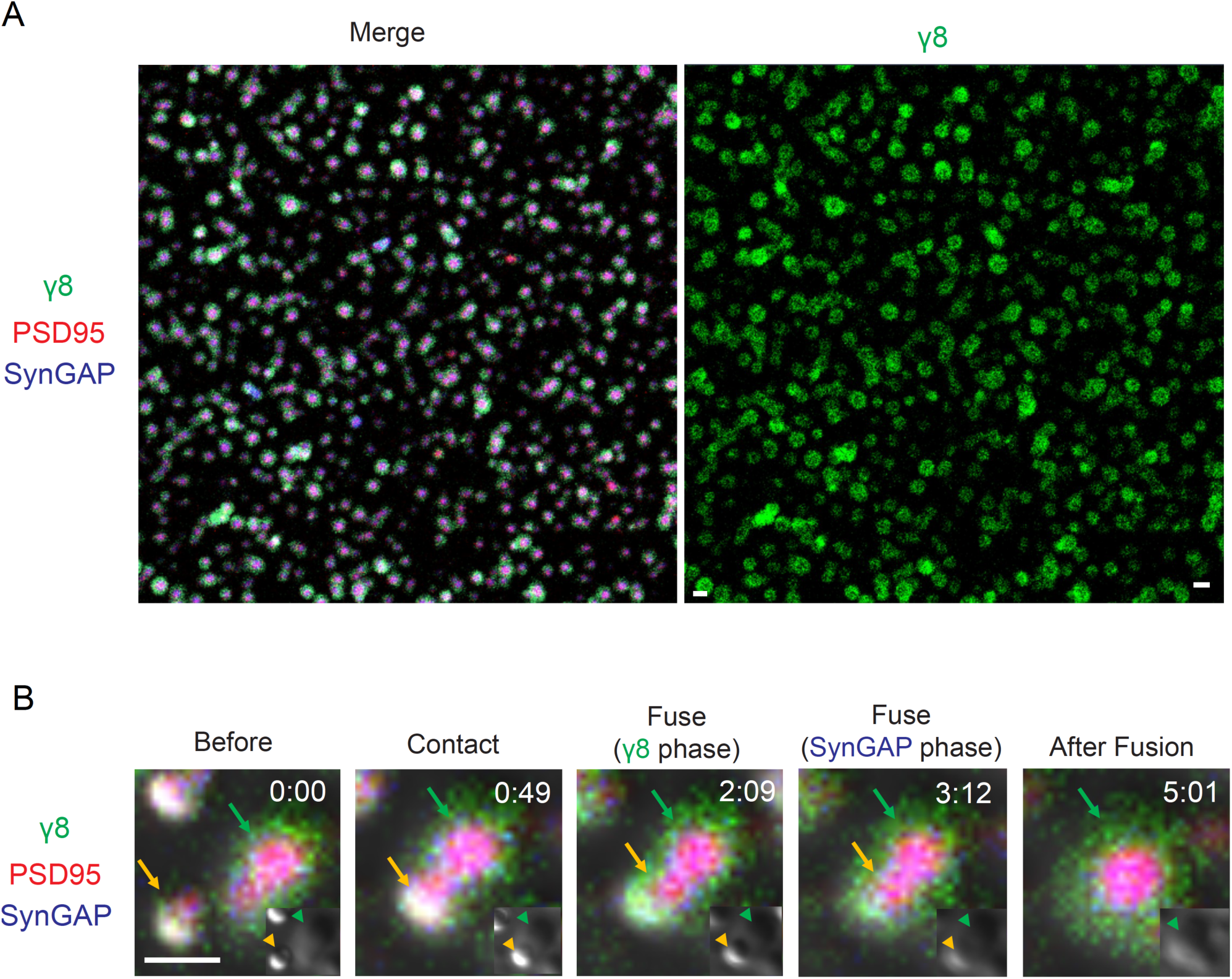
SynGAP-PSD95 and TARP-γ8-PSD95 show mutually exclusive phase-in-phase separation in droplets. (A) Additional representative wide field images of purified protein sedimentation assay under confocal microscopy. Purified proteins included TARP-γ8 (“γ8”) tagged with iFlour568 (Green), PSD95 tagged with iFlour633 (Red), and SynGAP tagged with iFlour488 (Blue). Scale bar 3 μm. γ8 localization was shown on the right panel to depict the uniformity of γ8 ring structure. Scale bar 3 μm. (B) Fusion events of γ8-PSD95-SynGAP protein droplets. Elapsed time (minutes: seconds) is shown at the top left of each panel. Green arrow: larger droplets at the bottom of the coverslip. Red arrow: smaller droplets precipitated and fused on larger droplets. Insets are DIC-only images.

**Figure S3.**
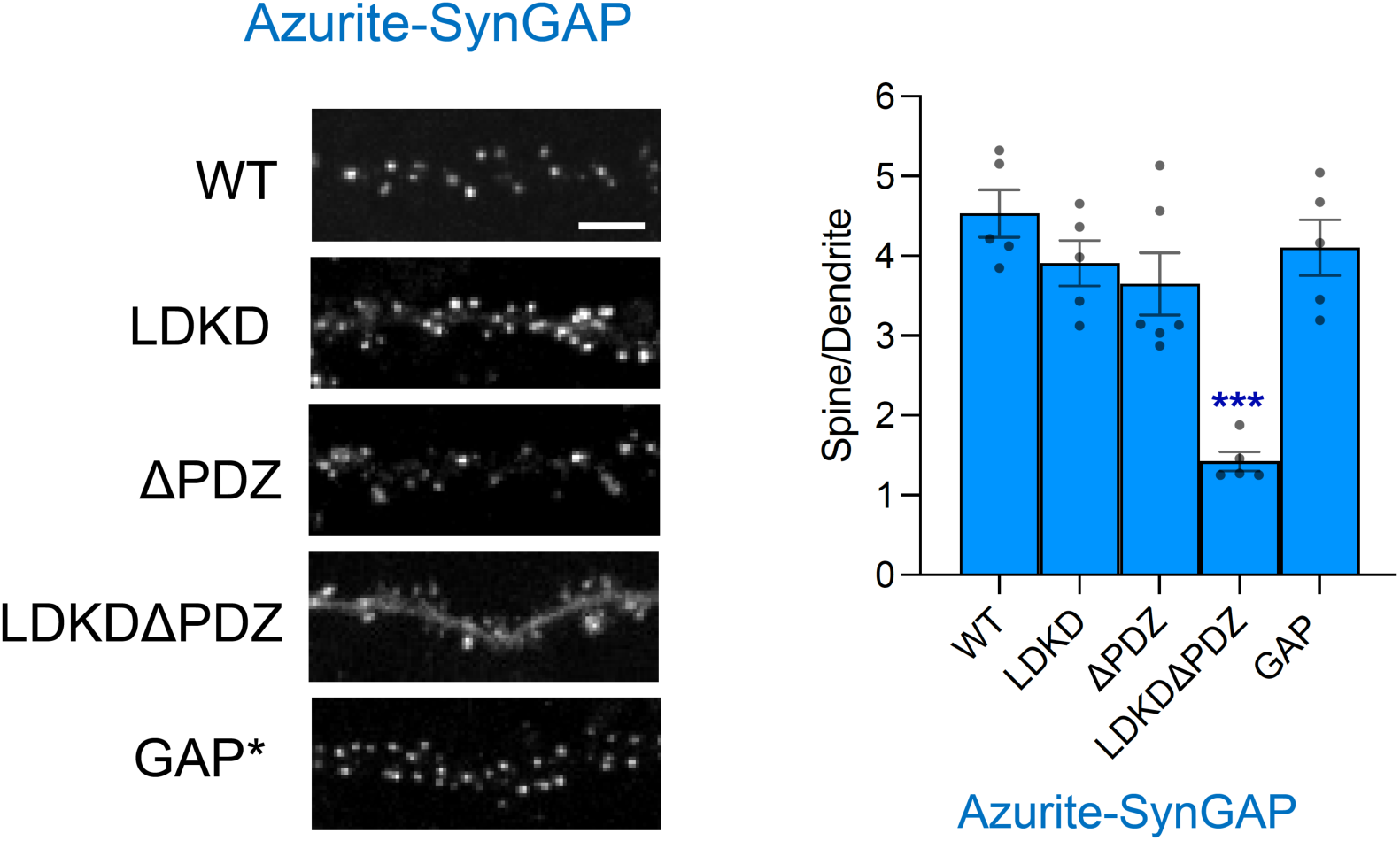
SynGAP mutations affecting PDZ-ligand or LLPS result in different SynGAP localization in rat hippocampal neurons. Confocal images of neurons expressing Azurite tagged various SynGAP mutations (PDZ ligand deficient or phase-separation deficient). Scale bar; 5 μm. SynGAP localization (signal from the Synaptic spine/signal from the dendritic shaft) was displayed on the right. One-way ANOVA with Tukey’s multiple comparisons test. Scale Bar; 5 μm. Error bars represent the S.E.M. *p<0.05, **p<0.01, ***p<0.001, ****p<0.0001, n.s. (not significant).

## References and Notes

1 Kessels, H. W. & Malinow, R. Synaptic AMPA receptor plasticity and behavior. Neuron 61, 340–350, doi:10.1016/j.neuron.2009.01.015 (2009).

2 Huganir, R. L. & Nicoll, R. A. AMPARs and synaptic plasticity: the last 25 years. Neuron 80, 704–717, doi:10.1016/j.neuron.2013.10.025 (2013).

3 Nicoll, R. A. A Brief History of Long-Term Potentiation. Neuron 93, 281–290, doi:10.1016/j.neuron.2016.12.015 (2017).

4 Shi, S. H., Hayashi, Y., Esteban, J. A. & Malinow, R. Subunit-Specific Rules Governing AMPA Receptor Trafficking to Synapses in Hippocampal Pyramidal Neurons. Cell 105, 331–343 (2001).

5 Lisman, J. & Raghavachari, S. A unified model of the presynaptic and postsynaptic changes during LTP at CA1 synapses. Science STKE 2006 (2006).

6 Opazo, P., Sainlos, M. & Choquet, D. Regulation of AMPA receptor surface diffusion by PSD-95 slots. Curr Opin Neurobiol 22, 453–460, doi:10.1016/j.conb.2011.10.010 (2012).

7 Araki, Y., Zeng, M., Zhang, M. & Huganir, R. L. Rapid dispersion of SynGAP from synaptic spines triggers AMPA receptor insertion and spine enlargement during LTP. Neuron 85, 173–189, doi:10.1016/j.neuron.2014.12.023 (2015).

8 Yang, Y., Tao-Cheng, J. H., Reese, T. S. & Dosemeci, A. SynGAP moves out of the core of the postsynaptic density upon depolarization. Neuroscience 192, 132–139, doi:10.1016/j.neuroscience.2011.06.061 (2011).

9 Walkup, W. G. et al. A model for regulation by SynGAP-alpha1 of binding of synaptic proteins to PDZ-domain ’Slots’ in the postsynaptic density. Elife 5, doi:10.7554/eLife.16813 (2016).

10 Mastro, T. L. et al. A sex difference in the response of the rodent postsynaptic density to synGAP haploinsufficiency. Elife 9, doi:10.7554/eLife.52656 (2020).

11 Kim, J. H., Liao, D., Lau, L. & Huganir, R. L. SynGAP: a Synaptic RasGAP that Associates with the PSD-95/SAP90 Protein Family. Neuron 20, 683–691 (1998).

12 Chen, H. J., Rojas-Soto, M., Oguni, A. & Kennedy, M. B. A synaptic Ras-GTPase activating protein (p135 SynGAP) inhibited by CaM kinase II. Neuron 20, 895–904, doi:10.1016/s0896-6273(00)80471-7 (1998).

13 Gamache, T. R., Araki, Y. & Huganir, R. L. Twenty Years of SynGAP Research: From Synapses to Cognition. J Neurosci 40, 1596–1605, doi:10.1523/JNEUROSCI.0420-19.2020 (2020).

14 Sheng, M. & Kim, E. The postsynaptic organization of synapses. Cold Spring Harb Perspect Biol 3, doi:10.1101/cshperspect.a005678 (2011).

15 Zhu, J. J., Qin, Y., Zhao, M., Aelst, L. V. & Malinow, R. Ras and Rap control AMPA receptor trafficking during synaptic plasticity. Cell 110, 443–455 (2002).

16 Komiyama, N. H. et al. SynGAP Regulates ERK-MAPK Signaling, Synaptic Plasticity, and Learning in the Complex with Postsynaptic Density 95 and NMDA Receptor. J. Neurosci. 22, 9721–9732 (2002).

17 Kim, J. H., Lee, H. K., Takamiya, K. & Huganir, R. L. The Role of Synaptic GTPase-Activating Protein in Neuronal Development and Synaptic Plasticity. J. Neurosci. 23, 1119–1124 (2003).

18 Guo, X. et al. Reduced expression of the NMDA receptor-interacting protein SynGAP causes behavioral abnormalities that model symptoms of Schizophrenia. Neuropsychopharmacology 34, 1659–1672, doi:10.1038/npp.2008.223 (2009).

19 Clement, J. P. et al. Pathogenic SYNGAP1 mutations impair cognitive development by disrupting maturation of dendritic spine synapses. Cell 151, 709–723, doi:10.1016/j.cell.2012.08.045 (2012).

20 Scheffzek, K. et al. The Ras-RasGAP Complex: Structural Basis for GTPase Activation and Its Loss in Oncogenic Ras Mutants. Science 277, 333–338 (1997).

21. Rumbaugh, G., Adams, J. P., Kim, J. H. & Huganir, R. L. SynGAP regulates synaptic strength and mitogen-activated protein kinases in cultured neurons. PNAS 103, 4344-4351 (2006).

22 Nakajima, R. et al. Comprehensive behavioral analysis of heterozygous Syngap1 knockout mice. Neuropsychopharmacol Rep 39, 223–237, doi:10.1002/npr2.12073 (2019).

23 Ozkan, E. D. et al. Reduced cognition in Syngap1 mutants is caused by isolated damage within developing forebrain excitatory neurons. Neuron 82, 1317–1333, doi:10.1016/j.neuron.2014.05.015 (2014).

24 Zeng, M. et al. Phase Transition in Postsynaptic Densities Underlies Formation of Synaptic Complexes and Synaptic Plasticity. Cell 166, 1163–1175 e1112, doi:10.1016/j.cell.2016.07.008 (2016).

25 Zeng, M. et al. Phase Separation-Mediated TARP/MAGUK Complex Condensation and AMPA Receptor Synaptic Transmission. Neuron 104, 529–543 e526, doi:10.1016/j.neuron.2019.08.001 (2019).

26 Wang, B. et al. Liquid-liquid phase separation in human health and diseases. Signal Transduct Target Ther 6, 290, doi:10.1038/s41392-021-00678-1 (2021).

27 Banjade, S. & Rosen, M. K. Phase transitions of multivalent proteins can promote clustering of membrane receptors. Elife 3, doi:10.7554/eLife.04123 (2014).

28 Zeng, M. et al. Reconstituted Postsynaptic Density as a Molecular Platform for Understanding Synapse Formation and Plasticity. Cell 174, 1172–1187 e1116, doi:10.1016/j.cell.2018.06.047 (2018).

29 Sumioka, A., Yan, D. & Tomita, S. TARP phosphorylation regulates synaptic AMPA receptors through lipid bilayers. Neuron 66, 755–767, doi:10.1016/j.neuron.2010.04.035 (2010).

30 Tidyman, W. E. & Rauen, K. A. Expansion of the RASopathies. Curr Genet Med Rep 4, 57–64, doi:10.1007/s40142-016-0100-7 (2016).

31 Cook, E. H., Masaki, J. T., Guter, S. J. & Najjar, F. Lovastatin Treatment of a Patient with a De Novo SYNGAP1 Protein Truncating Variant. J Child Adolesc Psychopharmacol 29, 321–322, doi:10.1089/cap.2018.0159 (2019).

32 Kluger, G., von Stulpnagel-Steinbeis, C., Arnold, S., Eschermann, K. & Hartlieb, T. Positive Short-Term Effect of Low-Dose Rosuvastatin in a Patient with SYNGAP1-Associated Epilepsy. Neuropediatrics 50, 266–267, doi:10.1055/s-0039-1681066 (2019).

33 Karczewski, K. J. et al. The mutational constraint spectrum quantified from variation in 141,456 humans. Nature 581, 434–443, doi:10.1038/s41586-020-2308-7 (2020).

34 Taliun, D. et al. Sequencing of 53,831 diverse genomes from the NHLBI TOPMed Program. Nature 590, 290–299, doi:10.1038/s41586-021-03205-y (2021).

35 Tadaka, S., et al. jMorp updates in 2020: large enhancement of multi-omics data resources on the general Japanese population. Nucleic Acids Res 49, D536–D544, doi:10.1093/nar/gkaa1034 (2021).

36 Vlaskamp, D. R. M. et al. SYNGAP1 encephalopathy: A distinctive generalized developmental and epileptic encephalopathy. Neurology 92, e96–e107, doi:10.1212/WNL.0000000000006729 (2019).

37 Mignot, C. et al. Genetic and neurodevelopmental spectrum of SYNGAP1- associated intellectual disability and epilepsy. J Med Genet 53, 511–522, doi:10.1136/jmedgenet-2015-103451 (2016).

38 Araki, Y. et al. SynGAP isoforms differentially regulate synaptic plasticity and dendritic development. Elife 9, doi:10.7554/eLife.56273 (2020).

